# Adaptive mitochondrial regulation of the proteasome

**DOI:** 10.1101/2020.04.07.026161

**Authors:** Thomas Meul, Korbinian Berschneider, Sabine Schmitt, Christoph H. Mayr, Laura F. Mattner, Herbert B. Schiller, Ayse Yazgili, Xinyuan Wang, Christina Lukas, Cornelia Prehn, Jerzy Adamski, Elisabeth Graf, Thomas Schwarzmayr, Fabiana Perocchi, Alexandra Kukat, Aleksandra Trifunovic, Laura Kremer, Holger Prokisch, Bastian Popper, Christine von Toerne, Stefanie M. Hauck, Hans Zischka, Silke Meiners

## Abstract

The proteasome is the main proteolytic system for targeted protein degradation in the cell. Its function is fine-tuned according to cellular needs. Regulation of proteasome function by mitochondrial metabolism, however, is unknown.

Here, we demonstrate that mitochondrial dysfunction reduces the assembly and activity of the 26S proteasome in the absence of oxidative stress. Impaired respiratory complex I function leads to metabolic reprogramming of the Krebs cycle and deficiency in aspartate. Aspartate supplementation activates assembly and activity of 26S proteasomes via transcriptional activation of the proteasome assembly factors p28 and Rpn6. This metabolic adaptation of 26S proteasome function involves sensing of aspartate via the mTORC1 pathway. Metformin treatment of primary human cells similarly reduced assembly and activity of 26S proteasome complexes, which was fully reversible and rescued by supplementation of aspartate or pyruvate. Of note, respiratory dysfunction conferred resistance towards the proteasome inhibitor Bortezomib.

Our study uncovers a fundamental novel mechanism of how mitochondrial metabolism adaptively adjusts protein degradation by the proteasome. It thus unravels unexpected consequences of defective mitochondrial metabolism in disease or drug-targeted mitochondrial reprogramming for proteasomal protein degradation in the cell. As metabolic inhibition of proteasome function can be alleviated by treatment with aspartate or pyruvate, our results also have therapeutic implications.

## Introduction

The ubiquitin-proteasome system is a central part of cellular protein homeostasis. It recycles amino acids for protein synthesis, maintains protein quality control, and controls the half-life of essential regulators of cell function [1–3]. Ubiquitin-dependent protein degradation is carried out by 26S proteasome complexes [4]. The 26S proteasome consists of a central catalytic core – the 20S proteasome – with one or two 19S regulators attached to the ends of the 20S forming single or double capped 26S proteasome complexes [5]. While the 20S proteasome core contains the three proteolytic sites with chymotrypsin-, caspase-, and trypsin-like activities for protein hydrolysis, the 19S regulator mediates ATP-dependent binding and unfolding of ubiquitinated substrates as well as recycling of ubiquitin moieties [5].

Mitochondria are the powerhouses of the cell, providing energy and redox equivalents via their respiratory chain [6]. Besides ATP production via the respiratory chain, mitochondria play key metabolic functions in the biosynthesis of nucleotides, lipids, and amino acids by generating essential intermediates via the Krebs cycle [7]. Dysfunction of the respiratory chain results in defective ATP production and metabolic reprogramming of the cell due to imbalanced redox equivalents [8–10].

Mitochondrial function is coupled to the ubiquitin-proteasome system [11–13]. Proteins of the inner and outer mitochondrial membrane are degraded in an ubiquitin-dependent manner by the 26S proteasome [14, 15]. Accumulation of misfolded or mistargeted mitochondrial proteins results in the adaptive activation of the ubiquitin-proteasome system to maintain mitochondrial and cellular proteostasis [16–19]. Conversely, 26S proteasome activity is reduced by defective ATP production after acute pharmacological inhibition of respiratory chain complexes [20] and 26S proteasome complexes disassemble upon excessive production of mitochondrial reactive oxygen species (ROS) [12,21,22]. On the other hand, defective proteasome function is compensated by adaptations of the mitochondrial metabolism [23].

We here demonstrate reversible fine-tuning of 26S proteasome activity by metabolic reprogramming of the cell. Specifically, we show that defective respiratory complex I function impairs the assembly of the 26S proteasome, which can be alleviated upon addition of aspartate or pyruvate. Assembly of 26S proteasome complexes depends on aspartate sensing via the mTORC1 pathway and subsequent transcriptional upregulation of specific proteasome assembly factors. Thus, our study uncovers a fundamentally new mechanism of adjusting protein degradation by the proteasome to the metabolic program of the cell, which has pathophysiological but also therapeutic consequences.

## Results

### Respiratory chain dysfunction reduces 26S proteasome activity and assembly

We set out to study metabolic regulation of proteasome activity in a model system of genetically impaired respiratory function without oxidative stress using mouse embryonic fibroblasts (MEF) of DNA mutator mice. These cells express a proof-reading deficient mutant of the mitochondrial DNA polymerase γ and accumulate random point mutations in their mitochondrial DNA (mtDNA) resulting in almost complete loss of respiratory chain activity (Suppl. Figures S1A-B). While mutator cells grew slower they did not show any morphological signs of stress (Suppl. Figures S1C-D). Experiments were performed with four different mutator and three distinct wildtype (WT) MEF cell lines to minimize clonal differences in the accumulation of mitochondrial DNA mutations [24].

The activity of all three proteolytic sites of the proteasome, namely the chymotrypsin-, caspase- and trypsin-like activities, was uniformly diminished in mutator cells when assaying specific substrate degradation (Figure 1A). We did not observe any differences in the expression of proteasomal subunits in WT and mutator cells (Figure 1B). Western blot analysis for 20S alpha and beta 5 subunits and for the 19S regulator subunits Rpn8 and TBP1 revealed similar expression levels. This finding was confirmed for RNA expression RNAseq (Figure 1C, Table S1) and RT-qPCR analysis for selected proteasome encoding genes (Suppl. Figure S1E). We next used native gels with substrate overlay analysis and immunoblotting to resolve the different active proteasome complexes in the cell. This method allows the discrimination into 20S proteasomes and single (26S) and double capped (30S) proteasome complexes. Cells with mitochondrial dysfunction showed reduced activity and amount of assembled single and double capped 26S proteasome complexes, while the level of free 20S proteasomes was slightly but not significantly increased (Figure 1D). The total amount of 20S complexes was marginally decreased in mutator compared to WT cells (Suppl. Figure S1F). This coincided with reduced levels of the 20S assembly factor Pomp in mutator cells (Suppl. Figure S1G).

**Figure 1:**
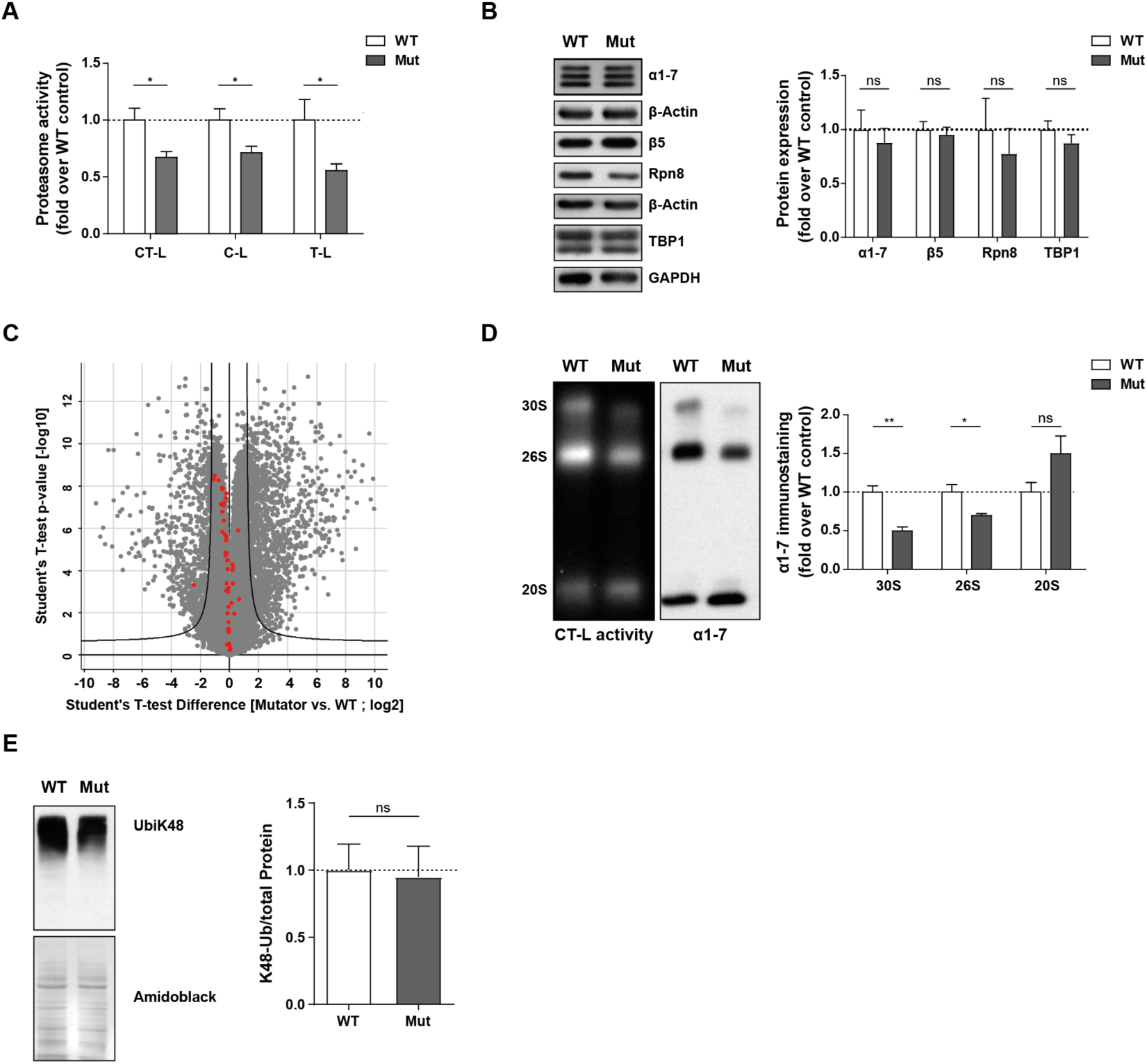
Respiratory chain dysfunction reduces 26S proteasome activity and assembly. Proteasome activity in total cell extracts of WT (n=3) and mutator (Mut) (n=4) cells as measured by cleavage of luminogenic substrates specific for the chymotrypsin-like (CT-L), caspase-like (C-L), or trypsin-like (T-L) active sites of the proteasome. Significance was determined using two-way ANOVA with Bonferroni multiple comparison test. (B) Representative Western blot analysis of 20S (α 1-7, β 5) and 19S regulator (Rpn8, Tbp-1) subunit expression in WT and mutator (Mut) cells. β-Actin or GAPDH were used as loading controls. Densitometric analysis shows mean±SEM from three WT and four mutator cell lines. Statistical test: Student’s unpaired t-test. (C) Volcano plot showing mRNA expression levels as identified by mRNA bulk sequencing of one WT (n=4 technical replicates) and one mutator (n=5 technical replicates) cell line using a 1 % FDR which is indicated by the black line. mRNA levels of proteasome subunits are shown in red. (D) Representative native gel analysis of active proteasome complexes and quantification thereof from native cell lysates from WT (n=3) and mutator (n=4) cells with chymotrypsin-like (CT-L) substrate overlay assay, immunoblotting for 20S α 1-7 subunits. The resolution of double capped (30S) and single capped 26S proteasomes (26S) is indicated together with the 20S proteasome complexes (20S). Bar graphs show mean±SEM relative to WT. Significance was determined using student’s unpaired t-test comparing WT vs. Mut cells. (E) Analysis of ubiquitinated proteins in WT and mutator cells. Representative Western blots of UbiK48-linked proteins. Bar graphs show Amido black normalized levels of UbiK48-linked proteins in WT (n=3) and mutator (Mut) cell lines (n=4) (mean±SEM). Significance was determined using student’s unpaired t-test.

26S proteasome deficiency in mutator cells might be caused by disassembly of 26S complexes into 20S catalytic core particles and 19S regulators due to diminished ATP or elevated ROS levels [20, 22]. Cellular ATP levels, however, were not reduced despite impaired mitochondrial ATP production (Suppl. Figure S1H) indicating maintenance of ATP production via elevated rates of glycolysis in mutator cells (Suppl. Figure S1I) as described before [25]. ROS levels were not increased in mutator cells as determined by cellular DCF and mitoSOX fluorescence (Suppl. Figure S1J) confirming previous results [25]. To also exclude effects of ROS micro-signaling, cells were treated with the antioxidant N-acetlycysteine (NAC). Proteasome activity in mutator cells was, however, not rescued by NAC treatment (Suppl. Figure S1K). These data indicate that reduced formation of 26S proteasome complexes in mutator cells does not involve ROS-mediated disassembly but rather reduced assembly of 26S proteasome complexes. Of note, diminished 26S proteasome activity did not alter the turnover of ubiquitinated proteins in mutator cells as levels of K48 polyubiquitinated proteins were similar to WT cells (Figure 1E). We also did not observe any signs of protein stress in our proteomic comparison of WT and mutator cells (Table S2) suggesting that reduced assembly of 26S proteasome complexes is part of an adaptive proteostatic cell response to mitochondrial dysfunction.

### Respiratory chain dysfunction results in aspartate deficiency

To elucidate the molecular mechanism involved in reduced 26S proteasome function in cells with mitochondrial dysfunction, we investigated the nature of mitochondrial alterations in mutator cells in detail. Cytochrome c staining revealed that neither the mitochondrial network nor their numbers were substantially altered in mutator compared to WT cells (Figure 2A, upper panel). We further purified intact mitochondria according to our recently developed protocol [26] and as schematically depicted in Suppl. Figure S2A. Mitochondria of mutator cells were structurally intact with only slight alterations in mitochondrial cristae structures as shown by electron microscopy of both cellular and isolated mitochondria (Figure 1A, lower panel, Suppl. Figure S2B). To confirm that the number of mitochondria is similar in WT and mutator cells the proportion of mitochondrial volume was quantified as described by Hacker and Lucocq (Suppl. Figure S2C) [27]. Proteomic analysis of isolated mitochondria from WT and mutator cells identified 714 mitochondrial proteins according to MitoCarta 2.0 [28] (Table S3). More than 90 % of these mitochondrial proteins were not regulated more than twofold. Using enrichment analysis on Uniprot keywords and Gene Ontology terms we observed severe downregulation of respiratory chain proteins (Figure 2B) (Tables S3+S4). These mapped mainly to complex I and complex IV as confirmed by immunoblot analysis (Suppl. Figure S2D). Complex I deficiency and the resulting lack of electron transfer resulted in impaired oxidation of NADH to NAD^+^ and pronounced accumulation of NADH in mutator cells (Figure 2C). Elevated NADH levels inhibit key enzymes of the Krebs cycle [29]. Together with the lack of NAD^+^ electron acceptors this hampers production of Krebs cycle intermediates that are required for the biosynthesis of macromolecules such as amino acids (Figure 2D) [8, 30]. Metabolomic quantification of amino acid levels revealed that the amount of aspartate was significantly decreased in mutator cells while overall levels of amino acids were not altered (Figure 2E, Suppl. Figure S2E). As aspartate is not supplemented in cell culture medium [31], cells strictly depend on the endogenous biosynthesis of aspartate for subsequent generation of amino acids, purines and pyrimidines [8,9,32]. Our analysis thus identifies deficiency of complex I, concomitant loss of electron acceptors, and reduced levels of aspartate as major features of metabolic reprogramming in mutator cells.

**Figure 2:**
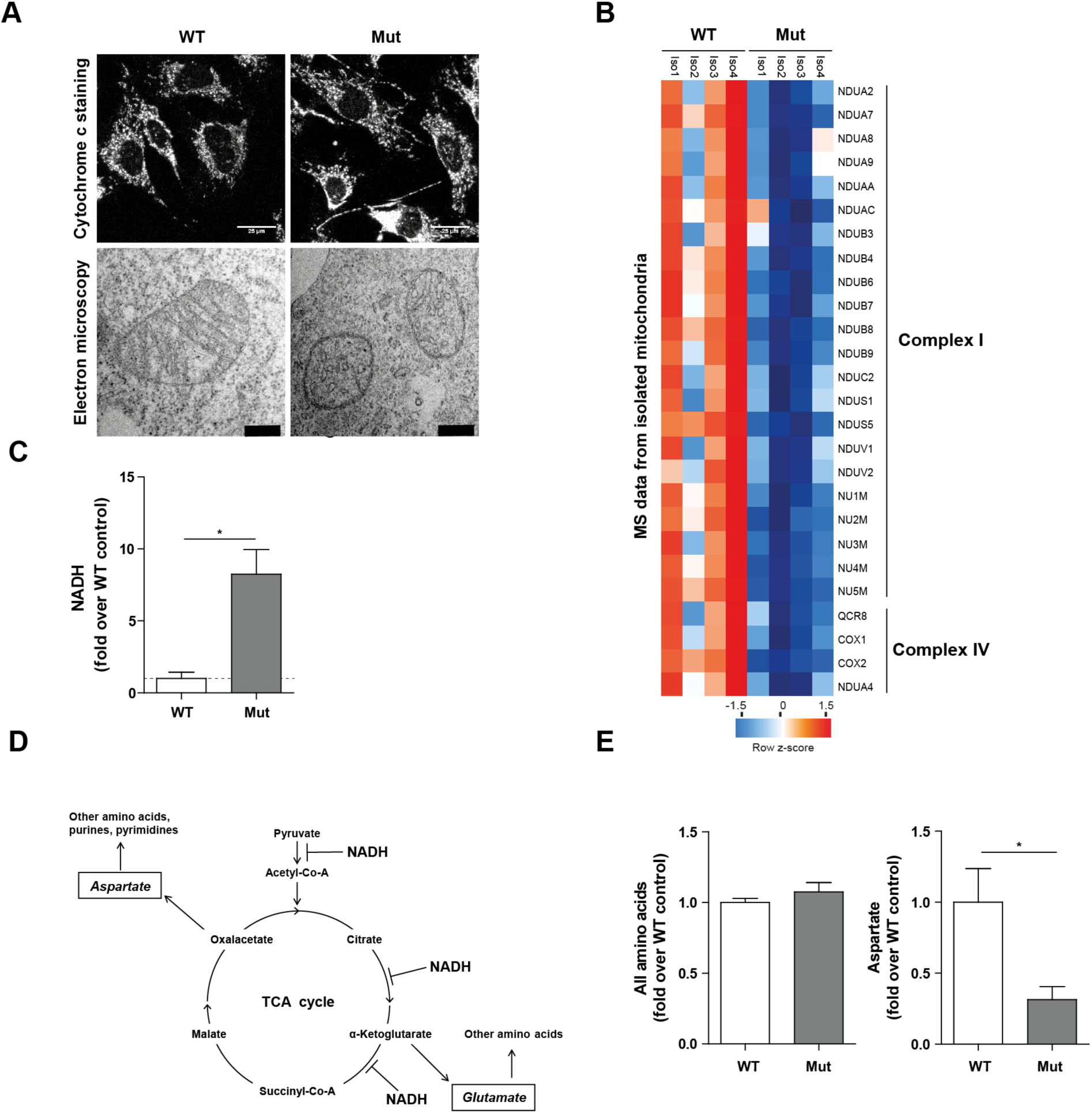
Respiratory chain dysfunction results in aspartate deficiency. Representative images of the mitochondrial network in WT (n=3) and mutator (n=4) cells visualized by staining with anti-cytochrome c antibody (upper panel). Scale bar: 25 µm. Lower panel shows representative electron microscopy images of mitochondria in a single WT and mutator cell line. Scale bar: 1 µm. (B) Heatmap representing z-scored relative protein mass-spectrometric intensities of significantly regulated respiratory chain complex subunits of mitochondria isolated from one mutator (4 technical replicates) and one WT (4 technical replicates) cell line. (C) NADH levels in WT (n=3) and mutator (n=4) cells. Bar graphs show mean±SEM relative to WT cells. Statistical test: student’s unpaired t-test. (D) Scheme of the TCA cycle emphasizing its role in amino acid synthesis and its inhibition by increased NADH levels. The amino acids aspartate and glutamate are directly synthesized from TCA cycle intermediates and serve as precursors for the synthesis of other amino acids and purines and pyrimidines. (E) Levels of all amino acids were determined by mass-spectrometry based targeted metabolomics in WT (n=3) and mutator (n=4) cells. 6 replicates were measured for each cell line and the respective values were normalized to the cell number of each cell line. Bar graphs show mean±SEM relative to WT controls. Significance was determined using student’s unpaired t-test.

### Supplementation of aspartate activates 26S proteasome activity

We next tested whether the observed aspartate deficiency in mutator cells contributes to the impairment of 26S proteasome activity. Aspartate supplementation for 72 hours fully activated the activity of the proteasome in mutator cells (Figure 3A). Aspartate treatment specifically increased the assembly and activity of mainly double capped 26S proteasomes (Figure 3B). Proteasome activation was first evident after 24 hours of aspartate treatment (Suppl. Figure S3A). Aspartate did not affect proteasome function in WT cells (Suppl. Figure S3B). Pyruvate - which serves as an electron acceptor at conditions of redox imbalance [8] - also fully restored single and double capped 26S proteasome assembly (Figure 3C) but did not affect proteasome activity in WT cells (Suppl. Figure S3C).

**Figure 3:**
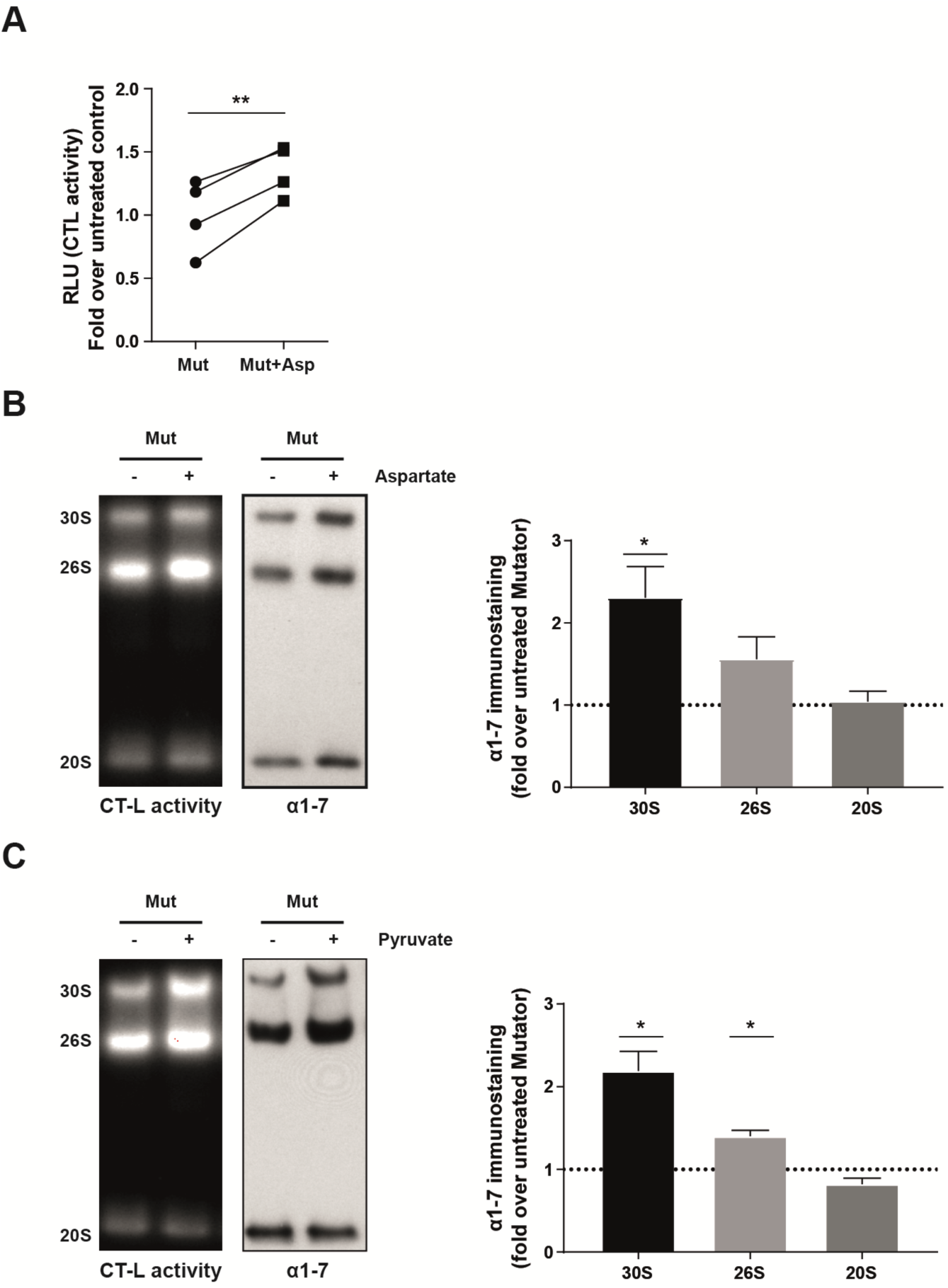
Supplementation of aspartate activates 26S proteasome activity. Proteasome activity in total cell extracts of mutator cells (n=4) treated with 10 mM aspartate for 72 h as determined by cleavage of luminogenic substrates specific for the chymotrypsin-like (CT-L) activity. Graph shows values of aspartate-treated cells that were normalized to the mean of untreated controls. Statistical test: student’s paired t-test. B) Representative native gel analysis of active proteasome complexes in cell lysates from mutator cells treated with 10 mM aspartate for 72 h. Chymotrypsin-like (CT-L) substrate overlay assay and immunoblotting for 20S α 1-7 subunits is shown. The resolution of double capped (30S) and single capped 26S proteasomes (26S) is indicated together with the 20S proteasome complexes (20S). Bar graphs represent mean±SEM of immunostaining signal relative to the respective untreated mutator cells (n=4 cell lines). Significance was determined using one sample t-test. C) Representative native gel analysis of active proteasome complexes in cell lysates from mutator cells treated with 1 mM pyruvate for 72 h. Chymotrypsin-like (CT-L) substrate overlay assay and immunoblotting for 20S α 1-7 subunits is shown. Bar graphs represent mean±SEM relative to the respective untreated mutator cells (n=3 cell lines). Significance was determined using one sample t-test.

### Aspartate supplementation induces assembly of 26S proteasomes

To investigate the underlying mechanism for metabolic activation of 26S proteasome assembly, we first analyzed whether aspartate supplementation affects expression of proteasome subunits. Overall expression of proteasomal subunits was not altered as determined by Western blotting for 20S alpha and the beta5 subunit (Figure 4A). In contrast, we noticed specific aspartate-mediated upregulation of several 26S proteasome assembly factors, i. e. the assembly chaperones p27 and p28 and the 19S subunit Rpn6, while S5b was not regulated by aspartate (Figure 4A). Aspartate induced RNA expression of the respective assembly factors (PSMD9, PSMD10 and PSMD11) already after 6 hours (Figure 4B). Silencing of p28 but not p27 counteracted assembly of 26S proteasomes only upon supplementation with aspartate as demonstrated by native gel and immunoblotting analysis (Figure 4C, Suppl. Figure S4A for controls). Partial silencing of Rpn6 similarly neutralized aspartate-induced 26S assembly and activation (Figure 4C, Suppl. Figure S4B). While p27 and p28 act as 19S regulatory particle assembly-chaperones (RACs) [33], Rpn6 is an essential part of the 19S regulator and serves as a molecular clamp that stabilizes the interaction with the 20S [34]. Accordingly, full depletion of Rpn6 is detrimental for the cell but an only minor increase in its expression level promotes assembly of 26S proteasome complexes [35, 36] while partial depletion counteracts 26S assembly [36]. Of note, the same assembly factors were downregulated in mutator compared to WT cells, while S5b - which is also known to inhibit 26S assembly [37, 38] - was upregulated (Suppl. Figure S4A). Our data thus demonstrate that metabolic regulation of 26S assembly factors is a key determinant in regulating the adaptive assembly of 26S proteasome complexes at conditions of mitochondrial dysfunction.

**Figure 4:**
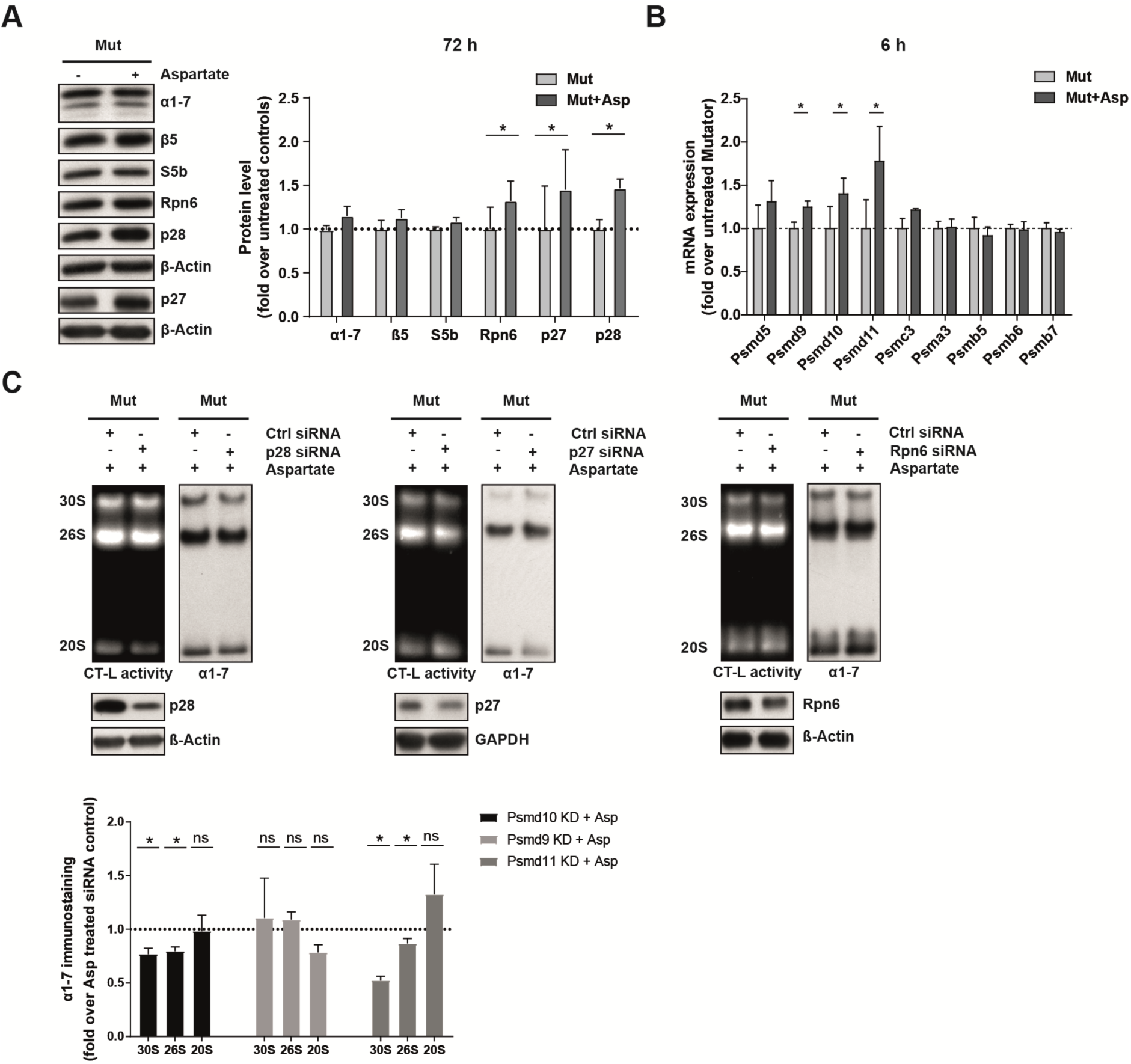
Aspartate supplementation induces assembly of 26S proteasomes. Representative Western blot analysis of 20S (α 1-7, β 5) and 19S (Rpn6, p27, p28, S5b) subunit expression in mutator cells treated with 10 mM aspartate for 72 h. β-Actin was used as a loading control. Densitometric analysis shows mean±SEM from four mutator cell lines. Significance was determined using student’s paired t-test. (B) RT-qPCR analysis of proteasome subunit mRNA expression in mutator (n=3) cells after 6 h of aspartate treatment. Data represent mean±SEM relative to untreated controls. Significance was determined using student’s paired t-test. (C) Representative native gel analysis of active proteasome complexes and quantification thereof in cell lysates from one mutator cell line upon p28 (n=4 independent experiments), p27 (n=3 independent experiments) and Rpn6 (n=5 independent experiments) silencing and aspartate treatment for 72 h with chymotrypsin-like (CT-L) substrate overlay assay and immunoblotting for 20S α 1-7. The resolution of double capped (30S) and single capped 26S proteasomes (26S) is indicated together with the 20S proteasome complexes (20S). Control cells were transfected with two different scrambled control siRNAs. Densitometric analysis shows mean±SEM of fold change over control from the respective replicate. Significance was determined using the one sample t-test. Knockdown was confirmed via immunostaining for p28, p27 and Rpn6. Only partial knockdown of Rpn6 was used to prevent cellular stress.

### Aspartate activates multiple cellular signaling pathways including mTOR signaling

We further dissected the mechanism by which aspartate activates 26S proteasome assembly in an unbiased phoshoproteome screen. Mutator cells were treated for four hours with or without aspartate. Phosphorylated peptides were enriched and identified by mass spec analysis according to a recently published protocol of the Mann lab [39]. We identified almost 10,000 phosphorylation sites with 233 phosphosites being significantly regulated by aspartate treatment. These mapped to 177 proteins (Table S5). About half of them were increased in abundance upon aspartate treatment (Suppl. Figure S5A). We identified numerous key regulators of the cell cycle, DNA replication, cytoskeleton, ribosome, transcription, and growth factor signaling pathways (Table S5). Proteasome subunits were not differentially phosphorylated by aspartate (Suppl. Figure S5B). Inspection of the kinase consensus motifs of the phosphorylated peptides identified several kinases that were activated by aspartate (Figure 5A, Table S6). Among these were several cell cycle related kinases such as CDK1, Aurora A, GSK3, ERK1, 2, and CDK5 besides other growth factor, cell cycle and metabolic signaling kinases (Figure 5A). This observation is well in line with the essential function of aspartate for proliferation [8, 9] and aspartate-mediated induction of proliferation in mutator cells (Suppl. Figure S5C). Other most predominantly activated kinases were the p70 ribosomal S6 kinase, MAPKAP1 and 2, and AKT kinases, which are all involved in the activation of protein synthesis via the mTOR pathway [40]. Activation of the mTOR pathway by aspartate was confirmed by enhanced phosphorylation of the p70 S6 kinase and the S6 ribosomal protein in Western blots (Figure 5B). In accordance with the activation of mTOR signaling, aspartate treatment for 72 hours increased protein synthesis in mutator cells (Figure 5C). Very similar, mTOR signaling and protein synthesis were found to be diminished in mutator compared to WT cells (Suppl. Figures S5D+E). Low doses of rapamycin (0.5 nM) effectively counteracted aspartate-induced activation of mTORC1 signaling (Suppl. Figure S5F). These data demonstrate a previously unrecognized regulation of protein synthesis by aspartate and indicate the presence of a specific aspartate sensor regulating mTORC1 function. Importantly, mTOR inhibition by rapamycin effectively prevented aspartate-induced upregulation of the proteasome assembly factors p28 and Rpn6 (Figure 5D) and activation of 26S proteasome assembly (Figure 5E). Rapamycin did not affect, however, proteasome activity and assembly in untreated mutator cells (Suppl. Figure S5G) Similar effects were observed upon silencing of raptor, a specific component of the mTORC1 complex [40] (Suppl. Figures S5H+I). These data demonstrate that aspartate activates the mTORC1 pathway, which - by a currently unknown pathway - induces the transcriptional activation of specific proteasome assembly factors to promote assembly of 26S proteasome complexes.

**Figure 5:**
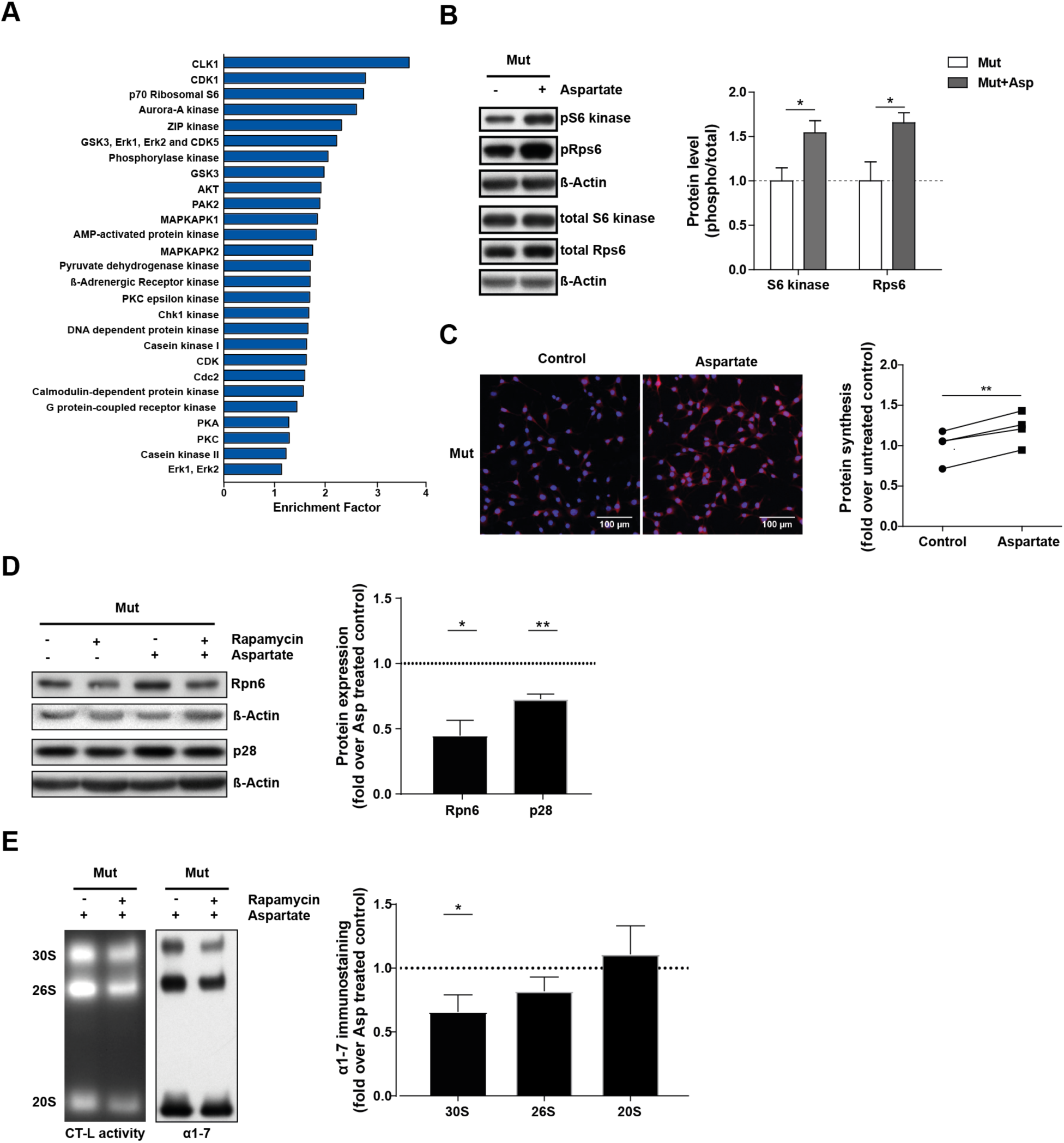
Aspartate activates multiple cellular signaling pathways including mTOR signaling. (A) Enrichment analysis of phospho proteomics data for kinases predicted to be activated upon aspartate treatment using fisher exact test (False discovery rate (FDR) > 0.02). (B) Analysis of mTORC1 signaling after treatment with 10 mM aspartate for 48 h in mutator cells. Representative Western blots of total and phosphorylated levels of p70 S6 kinase and S6 ribosomal protein (Rps6). β-Actin served as a loading control. Bar graphs show β-Actin normalized phosphoprotein levels related to β-Actin normalized total levels of the respective protein in mutator cell lines (n=4) (mean±SEM). Significance was determined using student’s unpaired t-test. (C) Cellular protein synthesis rate was determined using the puromycin analog OPP. Representative fluorescence images of nascent protein synthesis (red signal) and nucleic staining (blue signal) after supplementation of mutator cells (n=4) with 10 mM aspartate for 48 h are shown. Mutator controls were cultured for 48 h in normal medium containing no aspartate. Quantification of protein synthesis (Mean fluorescent intensity (MFI) of red signal) reveals differences in protein synthesis between the individual mutator cell lines with and without aspartate supplementation. Scale bar: 100 µm. Statistical test: student’s paired t-test. (D) Analysis of 26S proteasome assembly factor expression upon treatment with 0.5 nM rapamycin and 10 mM aspartate for 72 h in one mutator cell line. β-Actin was used as a loading control. Bar graphs show fold change of aspartate and rapamycin cotreatment over aspartate treated control. Significance was determined using the one sample t-test. (E) Representative native gel analysis of active proteasome complexes and quantification thereof in cell lysates from one mutator cell line upon treatment with 0.5 nM rapamycin and 10 mM aspartate for 72 h with chymotrypsin-like (CT-L) substrate overlay assay and immunoblotting for 20S α 1-7. Densitometric analysis shows mean±SEM of fold change of aspartate and rapamycin cotreatment over aspartate treated control from 3 independent experiments. Significance was determined using the one sample t-test.

### Defective complex I function drives metabolic adaptation of the proteasome in human cells

Our data obtained with the mutator cells strongly indicate that the defective electron transfer at complex I is the major driver for the observed metabolic regulation of 26S proteasome function. We thus tested whether the proteasome is metabolically regulated in cells with defined genetic complex I dysfunction. Skin fibroblasts were obtained from a patient with a hereditary mutation in the ND5 subunit of respiratory complex I [41]. These cells showed significantly diminished proteasome activity (Figure 6A), which we could attribute to reduced assembly and activity of 26S proteasome complexes (Figure 6B). Supplementation with aspartate activated assembly and activity of 26S proteasome complexes in ND5 mutant cells very similar to our results obtained with the mutator cells (Figure 6C). In a second approach, we applied pharmacological inhibition of complex I. For that, we used metformin - a well-known anti-diabetic drug which inhibits complex I activity in the absence of ROS production [8]. Of note, metformin treatment of primary healthy human skin fibroblasts for 72 hours significantly inhibited the assembly and activity of 26S proteasome complexes. This effect was fully restored by supplementation of either aspartate or pyruvate (Figure 6D). Similarly, metformin effectively inhibited 26S proteasome assembly in primary human lung fibroblasts, which was alleviated by treatment with aspartate or pyruvate (Figure 6E). Very similar results were obtained with WT MEFS (Suppl. Figure S6A).

**Figure 6:**
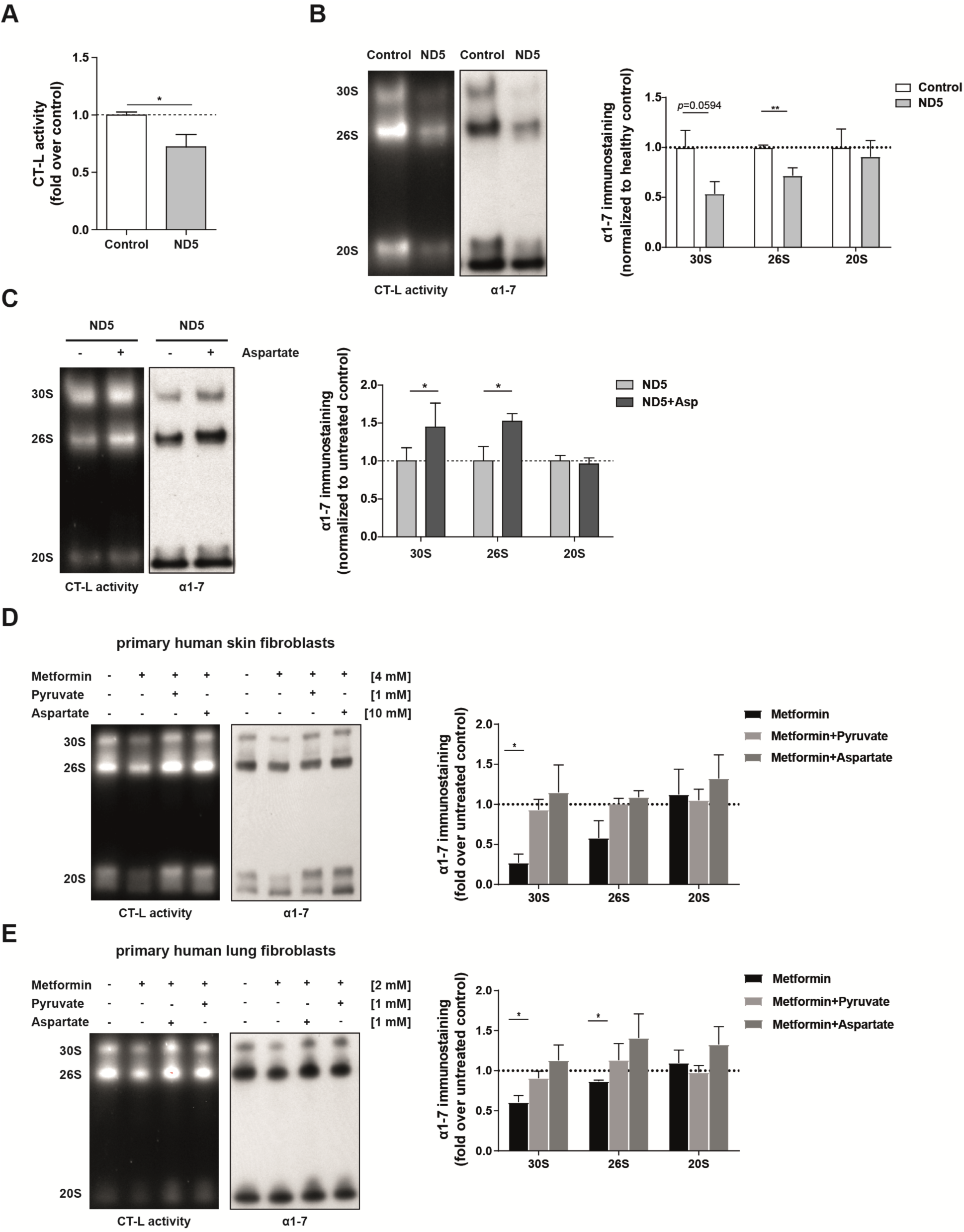
Defective complex I function drives metabolic adaptation of the proteasome in human cells. (A) Proteasome activity was measured in total cell extracts of ND5 mutant human fibroblasts and healthy controls in 5 different replicates from distinct cell passages using a luminogenic substrate specific for the chymotrypsin-like (CT-L) active site of the proteasome. Data represent mean±SEM relative to controls. Statistical test: student’s unpaired t-test. (B) Representative native gel analysis of active proteasome complexes in native cell lysates from human skin fibroblasts (healthy control and ND5 mutant) and quantification thereof with chymotrypsin-like (CT-L) substrate overlay assay and immunoblotting for 20S α 1-7. The resolution of double capped (30S) and single capped 26S proteasomes (26S) is indicated together with the 20S proteasome complexes (20S). Four independent experiments were performed. Bar graphs show mean±SEM relative to healthy control. Significance was determined using student’s unpaired t-test comparing healthy control vs. ND5 mutant cells. (C) ND5 mutant patient fibroblasts were treated with 10 mM aspartate for 48 h in six independent experiments. Activity and assembly of proteasome complexes was analyzed by native gel electrophoresis with CT-L substrate overlay assay and immunoblotting for 20S α 1-7 subunits. Densitometry shows mean±SEM values of aspartate-treated relative to untreated fibroblasts. Significance was determined using student’s unpaired t-test (D) Representative native gel analysis of active proteasome complexes in cell lysates from healthy primary human skin fibroblasts cotreated with 10 mM aspartate or 1 mM pyruvate together with 4 mM metformin for 72 h. Chymotrypsin-like (CT-L) substrate overlay assay and immunoblotting for 20S α 1-7 subunits is shown. Bar graphs represent mean±SEM relative to respective untreated skin fibroblasts (n=4 independent experiments). Significance was determined using one sample t-test. (E) Representative native gel analysis of active proteasome complexes in cell lysates from healthy primary human lung fibroblasts (phLF) cotreated with 1 mM aspartate or 1 mM pyruvate together with 2 mM metformin for 72 h. Chymotrypsin-like (CT-L) substrate overlay assay and immunoblotting for 20S α 1-7 subunits is shown. Bar graphs represent mean±SEM relative to respective untreated lung fibroblasts (n=4 independent experiments). Significance was determined using one sample t-test.

Our data thus reveal a reversible adjustment of 26S proteasome activity by aspartate or pyruvate under conditions of respiratory complex I dysfunction. They unravel a previous unknown cellular consequence of respiratory chain dysfunction for proteasomal protein degradation. Moreover, our findings suggest that metabolic inhibition of proteasome function can be alleviated by treatment with aspartate or pyruvate, which may have therapeutic implications.

### Respiratory chain dysfunction confers resistance to the proteasome inhibitor Bortezomib

To investigate the physiological relevance of reduced 26S proteasome activity and assembly in cells with respiratory dysfunction we tested the cellular response towards proteasome inhibition in mutator cells. WT and mutator cells were treated with increasing doses of the FDA approved proteasome inhibitor Bortezomib and cell viability was monitored. Of note, cells with mitochondrial dysfunction showed higher resistance towards Bortezomib-induced cell death (Figure 7A). Annexin V/PI staining confirmed reduced apoptotic cell death in mutator compared to WT cells with toxic doses of bortezomib (25 nM) (Figures 7B+C). This protective effect was specific for proteasome inhibition and not observed when cells were exposed to increasing concentrations of H_2_O_2_ (Figure 7D). From these observations we conclude that mitochondrial dysfunction contributes to resistance to proteasome inhibition possibly due to reduced activity of the 26S proteasome as recently suggested for bortezomib-resistant tumor cells [42–44].

**Figure 7:**
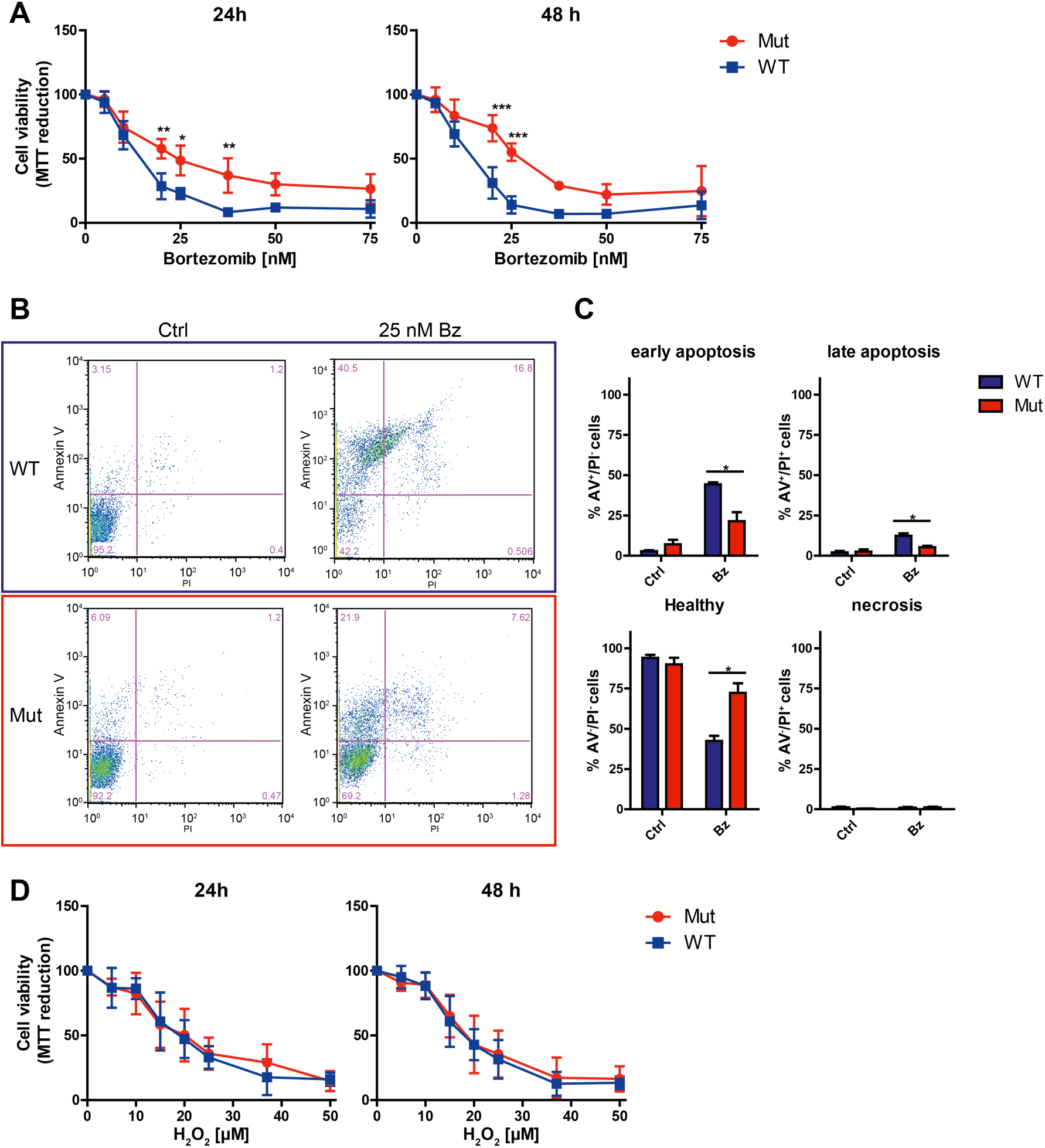
Respiratory chain dysfunction confers resistance to the proteasome inhibitor Bortezomib. (A) MTT cellular viability assay of WT (n=3) and mutator (n=4) MEFs treated with increasing doses of the proteasome inhibitor Bortezomib for 24 or 48 h. Values were normalized to the solvent-treated control (0 nM Bortezomib) and are displayed as mean+SEM. Significance was determined using two-way ANOVA with Bonferroni multiple comparison test. *:p< 0.05, **:p< 0.01, ***:p< 0.001. (B) Representative flow cytometry analysis of Annexin V and PI staining of WT and mutator MEFs treated with either solvent or 25 nM Bortezomib (Bz) for 24 h. (C) Quantification of Annexin V/Pi staining in WT (n=3) and Mut (n=4) cells. Cells were classified as healthy (Annexin V^-^/PI^-^), early apoptotic (Annexin V^+^/PI^-^), late apoptotic (Annexin V^+^/PI^+^), or necrotic (Annexin V^-^/PI^+^). Bar graphs show mean+SEM. Significance was determined using student’s t-test. *:p< 0.05. (D) Cellular viability of WT (n=3) and mutator (n=4) MEFs treated with increasing concentrations of H_2_O_2_ for 24 or 48 h. Values were normalized to the 0 µM H_2_O_2_-treated group and are displayed as mean+SEM.

## Discussion

In this study, we demonstrate that respiratory complex I dysfunction inhibits the activity of the ubiquitin-proteasome system by metabolic reprogramming. Impairment of 26S assembly is observed upon chronic impairment of mitochondrial function such as in mutator cells and ND5-mutant patient cells but also upon acute inhibition of complex I by metformin in the absence of oxidative stress. Importantly, inhibition can be fully reversed by supplementation of aspartate or pyruvate thus demonstrating adaptive metabolic fine-tuning of 26S proteasome function, which may have therapeutic implications.

It is well established that the 26S proteasome can be upregulated via concerted transcriptional induction of proteasomal gene expression under conditions of increased protein hydrolysis, protein stress, cell growth, and p53 signaling [45–48]. However, transcriptional regulation of proteasome function is tedious and time-consuming and not suitable for rapid and reversible adaptation to cellular needs [49, 50]. Rapid adjustment of 26S proteasome activity to growth signals can be achieved by posttranslational modifications of 19S or 20S subunits as e.g. elegantly shown for the 19S subunits Rpt3 [51, 52] and Rpn1 [53]. In addition, assembly chaperones regulate the formation of 26S proteasome complexes from the 20S catalytic core and 19S regulatory particles thus allowing timely adjustment of 26S proteasome activity to cellular demands [33, 50]. The regulatory particle assembly chaperones (RACs) S5b, p27, and p28 were found to be rapidly induced via ERK5 by inhibition of the mTOR pathway [54]. Similarly, phosphorylation of the constitutive 19S subunit Rpn6 by protein kinase A promoted assembly of 26S complexes upon mTOR inhibition [55, 56]. Such activation of 26S proteasome function under conditions of mTOR inhibition parallels that of autophagy induction to ensure cellular amino acids supply [40]. We here demonstrate an entirely different type of regulation: activation of the mTOR pathway by aspartate induced transcriptional upregulation of defined assembly factors, i.e. p27, p28, and Rpn6, but not 20S or other 19S subunits. This selective transcriptional activation makes it highly unlikely that Nrf1 is involved in this process [47]. Our phoshoproteomics analysis also excluded altered phosphorylation of Rpn6 and regulation of ERK5 to be involved in this process. Importantly, the two assembly factors p28 and Rpn6 were required for the induction of 26S proteasome assembly by supplementation of aspartate. We also demonstrate that this regulation is reversible as aspartate or pyruvate supplementation was sufficient to fully restore defective 26S proteasome assembly upon metformin treatment. Nucleoside addition, however, did not rescue proteasome assembly. These data suggest that nucleotide deficiency is not rate limiting but rather the apparent lack of electron acceptors. The adaptive nature of this regulation is also supported by the fact that the cell does not display any signs of protein stress such as accumulation of polyubiquitinated proteins or folding chaperones at conditions of reduced 26S proteasome activity. Of note, reversible adjustment of 26S proteasome assembly was evident in different cell types, i.e. mouse embryonic fibroblasts and primary human skin or lung fibroblasts, indicating a robust and conserved mechanism of metabolic fine-tuning of proteasome function. These findings add an important aspect of proteasome regulation and fit to the emerging concepts of fine-tuning of proteasome activity and also ribosome function according to cellular needs [49,50,57].

Our findings establish a fundamental novel interaction between mitochondrial function and the ubiquitin-proteasome system. Specifically, impaired activity of respiratory complex I and subsequent reprogramming of the Krebs cycle with deficiency of aspartate and electron acceptors results in reduced 26S proteasome activity. This novel interaction is of significant relevance to aging and disease: Both, aging and diseases of proteostasis imbalance such as neurodegenerative and cardiac diseases are characterized by respiratory chain dysfunction and impaired proteasome activity [58–63]. It is tempting to speculate that glycolytic reprogramming of tumor cells might contribute to diminished protein synthesis and 26S proteasome activity to confer resistance towards proteasome inhibitor treatment. Impaired 26S activity has previously been established as a prominent feature of proteasome inhibitor resistance [42–44]. Moreover, recent work also supports the link of altered mitochondrial function with proteasome inhibitor resistance [23]. Activating 26S proteasome assembly and activity by supplementation of aspartate or pyruvate might provide a therapeutic concept to counteract imbalanced proteostasis in disease and upon drug treatment.

## Author contributions

T.M., K.B., S.S., C.L., A.Y., X.W., C.P., L.K., C.vT., B.P. performed the experiments. T.M., K.B., S.S., C.P., C.H.M., H.B.S., J.A, L.K., H.P., F.P., O.E., C.vT., S.M.H., H.Z., S.M. analyzed and discussed the data. H.P., A.K., A.T. provided cells and animals. T.M., C.P., S.S., C.H.M. prepared the figures. T.M., S.S., H.Z., S.M. wrote the manuscript.

## Acknowledgement

The authors declare that they have no conflicts of interests. Korbinian Berschneider was supported by the CPC Graduate School “Lung biology and disease”. We are grateful for the support by Jennifer Wettmarshausen, and for the excellent technical assistance of Carola Eberhagen.

## STAR Methods

### Cell culture

Mouse embryonic fibroblasts from mtDNA mutator mice were generated as previously described [64]. Wildtype (WT) and mutator MEFs were cultured in DEMEM High Glucose (4.5 g/l) medium (without Sodium pyruvate) (ThermoFisher Scientific, Rockford, USA) supplemented with 10 % FBS, 1 % penicilin/streptomycin at 37 °C and 5 % CO_2_. MEFs isolated from four individual mtDNA mutator and three individual WT mice were used as different biological replicates [25]. Primary skin fibroblasts were cultured in DEMEM High Glucose (4.5 g/l) medium (without sodium pyruvate) (ThermoFisher Scientific, Rockford, USA) supplemented with 10 % FBS, 1 % penicilin/streptomycin and 200 µM uridine at 37 °C and 5 % CO_2_. Cells were grown to approx. 70 % confluency before passaging them into new flasks or using them for experiments. At least three different passages of the individual patients’ lines were used as biological replicates. Primary human lung fibroblasts were cultured in DEMEM High Glucose (4.5 g/l) medium (without sodium pyruvate) (ThermoFisher Scientific, Rockford, USA) supplemented with 10 % FBS, 1 % penicilin/streptomycin, 2 mM glutamine (ThermoFisher), 2 ng/ml β-FGF (ThermoFisher), 0.5 ng/ml EGF (Sigma-Aldrich) and 5 µg/ml insulin (ThermoFisher).

### Protein extraction

To preserve proteasome activity native protein lysates were prepared. Cell pellets were resuspended in TSDG buffer (10 mM Tris/HCl, 25 mM KCl, 1.1 mM MgCl2, 0.1 mM EDTA, 1 mM DTT, 1 mM NaN3, 10% glycerol, pH 7) containing complete protease inhibitor (Roche Diagnostics, Basel, Switzerland) and phosphatase inhibitor (PhosStop, Roche Diagnostics). Cell suspensions were subjected to seven cycles of freezing in liquid nitrogen and thawing at room temperature. Cell debris was removed by centrifugation and protein concentration in the supernatant was assessed using the Pierce BCA protein assay (ThermoFisher Scientific).

### Proteasome activity assays

In order to determine the activity of the three 20S proteasome active sites namely chymotrypsin-like (CT-L), caspase-like (C-L) and trypsin-like (T-L) the Proteasome-Glo™ Assay (Promega, Fitchburg, USA) was applied according to the manufacturer’s instructions. Here, 1 μg protein (TSDG lysates) per sample was first diluted with TSDG buffer to obtain equal volumes of TSDG buffer in each sample and then adjusted to a final volume of 20 μl with water. The prepared dilutions were transferred to white flat bottom 96-well plates and mixed with 20 μl of the respective active site specific substrate Suc-LLVY-aminoluciferin (CT-L specific), Z-nLPnLD-aminoluciferin (C-L specific) and Z-LRR-aminoluciferin (T-L specific). The different peptide substrates are cleaved by the respective proteasome active site and the released aminoluciferin serves as a substrate for the luciferase contained in the reaction buffer to generate a luminescent signal. This light signal was measured every 5 min for 1 h during the reaction in a Tristar LB 941 plate reader (Berthold Technologies, Bad Wildbad, Germany). For each substrate the samples were measured in technical triplicates.

### Western blot analysis

For Western blot analysis, 15 µg of protein lysates were mixed with 6x Laemmli buffer (300 mM Tris, 50 % Glycerol, 6 % SDS, 0.01 % Bromphenol blue, 600 mM DTT) and incubated at 95 °C for 5 min. After the incubation, samples were subjected to electrophoresis on 10 - 15 % SDS PAGE gels and blotted onto polyvinylidene difluoride (PVDF) membranes. Electrophoretic separation was performed at 90 – 130 V at room temperature and transfer to PVDF membranes was performed at 250 mA for 90 min at 4 °C. Membranes were blocked using Rotiblock (Roth, Karlsruhe, Germany) and treated with primary antibodies overnight at 4 °C and subsequently with secondary antibodies (1:40000) for 1 h at room temperature. Antibodies used were: Anti-α1-7 (1:1.000, Abcam, Cambridge, UK), Anti-Rpn6 (1:1000, Novus Biologicals, Littleton, CO, USA), Anti-β5 (1:1000, Santa Cruz Biotechnology, Dallas, TX, USA), Anti-Tbp1 (1:1000, BETHYL), Rpn8 (1:1000, Abcam), Anti-S5b (1:1000, Abcam), Anti-p27 (1:1000, Abcam), Anti-p28 (1:1000, Abcam), Anti-S6 kinase (1:2000, Cell Signaling), Anti-phospho p70 S6 kinase (1:2000, Cell Signaling), Anti-S6 ribosomal protein (1:2000, Cell Signaling), Anti-phospho S6 ribosomal protein (1:2000, Cell Signaling), Anti-Raptor (1:1000, Cell signaling), Anti-K48-Ubiquitin (1:3000, Merck-Millipore), HRP-conjugated anti-β-Actin (1:40.000, Sigma-Aldrich, Taufkirchen, Germany), HRP-conjugated anti-GAPDH (1:20.000, Cell signaling), Anti-OxPhos Rodent WB Antibody Cocktail (Cat No 45-8099 Thermo Fisher Scientific, Waltham, USA).

### Native gel analysis

15 μg protein of TSDG lysates were diluted with TSDG buffer to the highest sample volume. The diluted lysates were mixed with 5x native loading buffer (250 mM Tris, 50 % Glycerol, 0.01 % Bromphenol blue) and were loaded onto commercially available 3-8 % gradient NuPAGE Novex Tris-acetate gels (Life Technologies, Darmstadt, Germany). Native proteasome complexes were separated at 150 V for 4 h at 4 °C. After native gel electrophoresis the CT-L activity of the separated native proteasome complexes was determined by an in-gel substrate overlay proteasome activity assay. For this purpose gels were incubated for 30 min at 37 °C in a buffer containing 50 mM Tris, 1 mM ATP, 10 mM MgCl2, 1 mM DTT and 0.05 mM Suc-LLVY-AMC (Bachem, Bubendorf, Switzerland). Fluorescence generated by the CT-L specific cleavage of the fluorogenic Suc-LLVY-AMC peptide substrate in the active proteasome was detected within the gel at an excitation wavelength of 380 nm and emission wavelength of 460 nm using the ChemiDoc XRS+ system (BioRad, Hercules, USA). Finally, the gels were incubated for 15 min in a solubilization buffer (66 mM Na_2_CO_3_, 2 % SDS, 1.5 % β-mercaptoethanol) at room temperature in order to facilitate the subsequent blotting of the proteins using the previously described Western blot technique.

### Quantitative real-time RT-PCR

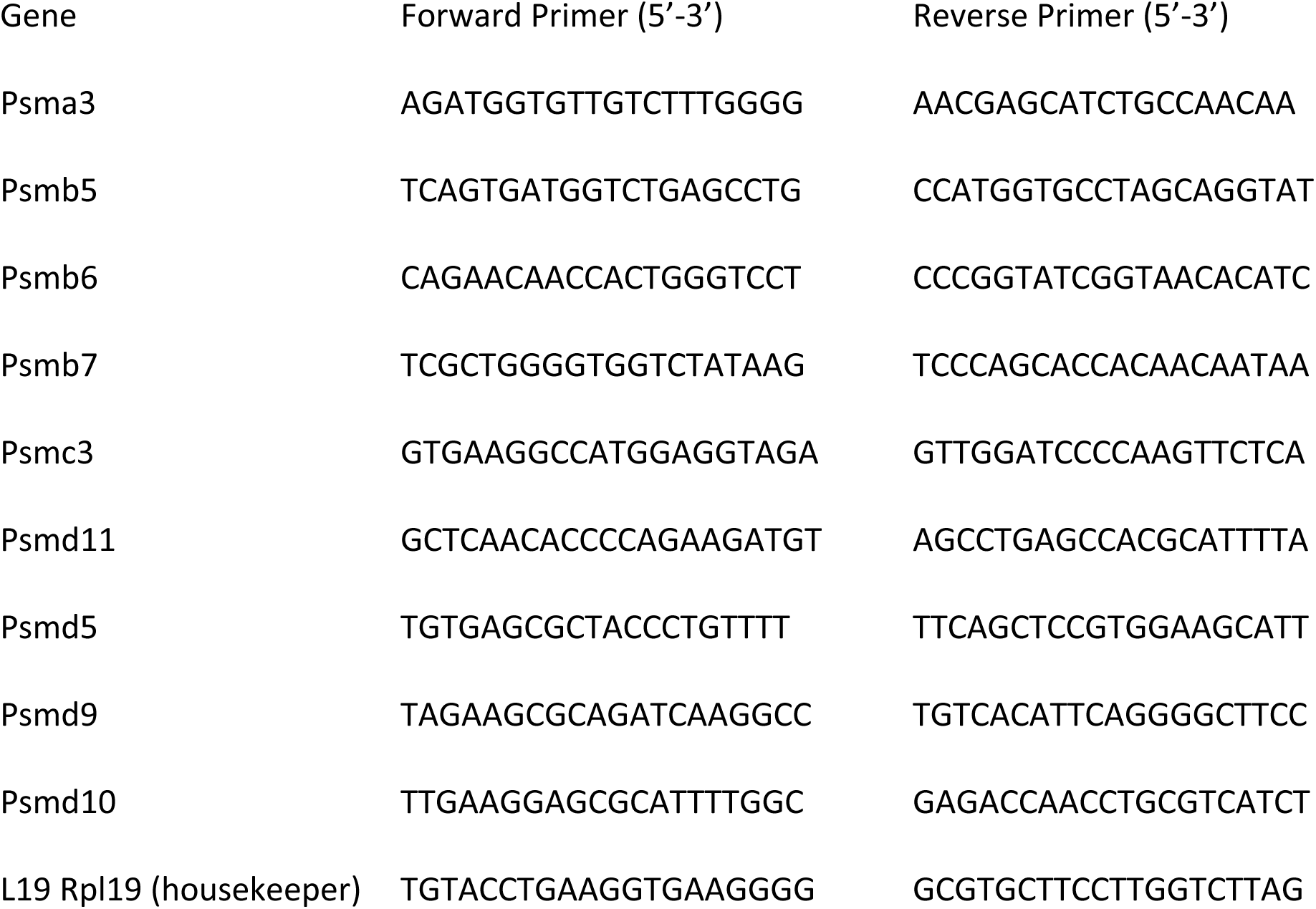

Total RNA isolation was performed by phenol/chloroform extraction using the Roti-Quick-Kit (Roth, Karlsruhe, Germany). 1 μg RNA per sample was transcribed into cDNA by M-MLV reverse transcriptase (Sigma Aldrich, Taufkirchen, Germany). The obtained cDNA was applied for quantitative RT-PCR with the SYBR Green LC480 System (Roche Diagnostic, Mannheim, Germany) using the respective forward and reverse primers for the genes of interest (Psma3, Psmb5, Psmb6, Psmb7, Psmc3, Psmd11, Psmd5, Psmd9, Psmd10) at a concentration of 2 pM. All samples were measured in technical duplicates.

### Measurement of ATP levels

Cellular ATP levels were determined using the CellTiter-Glo assay kit (Promega, Madison, WI, USA). Cells were seeded in 6-well plates and cultured for 24 h. After the treatment cells were harvested with trypsin and 40.000 cells/well were transferred to a white flat bottom 96-well plate. CellTiter-Glo reagent was added and the luminescent signal was measured after 3 min at a time for 30 min in total in a Tristar LB 941 plate reader (Berthold Technologies).

### Analysis of reactive oxygen species (ROS) in cells

Mitochondrial superoxide generation was analyzed using the MitoSOX® Red probe (Life Technologies) and overall cellular ROS generation was analyzed by H2DCFDA (ThermoFisher Scientific) fluorescence. For fluorescence analysis cells were seeded in 6-well plates and grown for 24 h. Cells were then stained for 30 minutes in medium containing 5 µM MitoSOX® Red or 25 µM H2DCFDA, respectively. Cells were washed with PBS, trypsinized, and resuspended in 500 µl FACS Buffer (PBS, 2 % FBS, 20 µM EDTA). Samples were then analyzed by flow cytometry using a BD LSRII Flow Cytometer (BD Biosciences, Heidelberg, Germany) and mean fluorescence intensity was measured.

### Cellular oxygen consumption

Cellular oxygen consumption rates were determined using the Seahorse XF Analyzer as described before [65].

### Mitochondrial cytochrome c staining

To analyze the mitochondrial network in WT and mutator MEFs, cells were seeded on 15 mm glass cover slips and cultured to 50 % confluency. Cells were then fixed on the cover slips using 4 % paraformadehyde for 10 min. Permeabilization was performed with 0.1 % Triton-X in PBS for 15 min and unspecific binding sites were blocked with Roti-Immunoblock (Roth, Karlsruhe, Germany) for 1 h at room temperature. Cells were then incubated with anti-cytochrome c antibody (1:1000, BD Bioscience, San Jose, CA, USA) for 2 h at room temperature. The secondary Alexa Fluor® (AF)-488-coupled antibody (1:750, Life Technologies) was added for 1 h at room temperature. After 2 washing steps in PBS cells were stained with 4’-6-Diamidin-2-phenylindol (DAPI) (Sigma-Aldrich) for 5 min. For confocal laser microscopy (Zeiss LSM710, Oberkochen, Germany) cells were mounted on microscopic slides using DAKO mounting medium.

### Isolation of mitochondria

Mitochondria from cultured cells were isolated as previously described [66]. Briefly, cells were resuspended in isolation buffer (300 mM sucrose, 5 mM TES, 200 µM EGTA, pH 7.2, without BSA) to a concentration of 5-7×10^6^ cells per ml and pumped 4 times through a clearance of 6 µm (flow rate 700 µl/min). The homogenate was centrifuged at 800 x g (4°C) to remove nuclei and cell debris and mitochondria were pelleted at 10000 x g. For purification, mitochondria were loaded on a Nycodenz^®^ (Axis Shield PoC AS, Oslo, Norway) density gradient (24%/18%) and centrifuged at 30.000 rpm for 15 min at 4°C in a Beckman ultracentrifuge (rotor SW 55.Ti). Mitochondria were collected at the 24%/18% interphase and washed once in isolation buffer without BSA (9000 x g, 10 min at 4°C).

### Electron microscopy

Electron microscopy of cells and there from isolated mitochondria was done as previously described [67]. Samples were fixed with 2.5% glutaraldehyde in 0.1M sodium cacodylate buffer, pH 7.4 (Electron Microscopy Sciences, Hatfield, USA) for 24 h at minimum. Thereafter glutaraldehyde was removed and samples were washed three times with 0.1M sodium cacodylate buffer, pH 7.4. Postfixation and prestaining was done for 45 to 60 min with 1% osmium tetroxide (10ml 4% osmium tetroxide (Electron Microscopy Sciences, cat no. 19190, Hatfield, USA), 10ml ddH2O, 10ml 3.4% sodium chloride and 10ml 4.46% potassium dichromate (pH adjusted to 7.2 with KOH (Sigma Aldrich)). Samples were washed three times with ddH_2_O and dehydrated with an ascending ethanol series (15min with 30%, 50%, 70%, 90% and 96%, respectively and two times 10min with 100%) and propylene oxide (two times 30min, Serva Electrophoresis GmbH, Heidelberg, Germany). Subsequently, samples were embedded in Epon (3.61M glycidether 100, (Serva Electrophoresis GmbH), 1.83M methylnadicanhydride (Serva Electrophoresis GmbH), 0.92M dodecenylsuccinic anhydride (Serva Electrophoresis GmbH), 5.53mM 2,4,6-Tris(dimethylaminomethyl)phenol (Serva Electrophoresis GmbH)). Ultrathin sections were automatically stained with UranyLess EM Stain (Electron Microscopy Sciences) and 3% lead citrate (Leica, Wetzlar, Germany) using the contrasting system Leica EM AC20 (Leica, Wetzlar, Germany) and examined with a JEOL-1200 EXII transmission electron microscope at 80 kV (JEOL GmbH, Freising, Germany). Images were taken using a digital camera (KeenViewII, Olympus, Germany) and processed with the iTEM software package (anlySISFive ; Olympus, Germany).

### Measurement of cellular NADH levels

Cellular NADH levels in WT and mutator MEFs were determined using the NAD/NADH-Glo assay kit (Promega) according to the manufacturer’s instructions. Cells were trypsinized and washed with PBS. 40000 cells per sample (triplicates) dissolved in 50 µl PBS were transferred in a 96-well plate. Cell lysis was performed using 50 µl 0.2 M NaOH with 1 % Dodecyltrimethylammonium bromide (DTAB). Cells were incubated at 60 °C for 15 min to deplete NAD^+^ from the lysates. 50 µl 0.25 M Tris, 0.2 M HCl solution was added to the wells. 20 µl of each well were transferred into a white flat bottom 96-well plate and mixed with 20 µl NAD/NADH Glo detection reagent. Obtained luminescence was measured in a Tristar LB 941 plate reader.

### Cell viability assay (MTT)

Cellular viability was measured using the 2,5-diphenyltetrazolium bromide (MTT) assay. 6000 cells/well were seeded into 96-well plates with each sample in at least three technical replicates and treated with increasing doses of the proteasome inhibitor Bortezomib (Sigma-Aldrich, St. Louis, MO). Cells were grown for 24 or 48 h and medium was changed to 100 µl treatment or control medium as indicated in the figures. After treatment, 20 µl of freshly prepared thiazolyl blue tetrazolium bromide solution (5 mg/ml in PBS) (Sigma-Aldrich, St. Louis, MO) was added to each well and cells were incubated for 30 min at 37°C. The supernatant was aspirated, and blue crystals were dissolved in isopropanol + 0.1 % Triton X-100. Absorbance was measured at 570 nm using a Tristar LB 941 plate reader.

### Live/dead staining with Annexin V/PI

Induction of apoptosis or necrosis was analyzed by staining cells with an Annexin V antibody and Propidium Iodide (PI). Cells were seeded into 6-well plates and grown for 24 h. Cells were then treated with Bortezomib for 24 h. After the treatment, cells were trypsinized, washed twice with PBS and resuspended in 100 µl Annexin V binding buffer. 5 µl anti-Annexin V-FITC (BD Biosciences, San Jose, CA) and 10 µl PI staining solution were added and cells were incubated for 15 min at room temperature in the dark. After the incubation time, 400 µl Annexin V binding buffer were added and fluorescence was measured by flow cytometry analysis using the Becton Dickinson LSRII and analyzed using FlowJo software (version 7.6.5).

### Metabolomic analysis

Wildtype and mutator MEFs were seeded in 6 well plates for a final cell number of around 1 million cells after 24 h. 6 wells per cell line were used for metabolomics analysis and 6 additional wells were counted to determine the final cell number of each cell line. Cells were first washed two times with PBS and then overlaid with 300 µl dry ice cold methanol. Cells of each well were harvested by scraping and transferred into a 0.5 mL PP-Sarstedt Micro tube (Sarstedt AG & Co, Nümbrecht, Germany) and frozen immediately on dry ice. Samples were stored at −80 °C until metabolomics analysis. Targeted metabolomics analysis was performed at the Helmholtz Zentrum München, Institute of Experimental Genetics, Genome Analysis Center in Neuherberg, Germany. Metabolites were quantified using the Absolute*IDQ*^TM^ Kit p180 (BIOCRATES Life Sciences AG, Innsbruck, Austria) and LC-ESI-MS/MS and FIA-ESI-MS/MS measurements as described previously [68]. The method of Absolute*IDQ*^TM^ p180 Kit has been proven to be in conformance with the EMEA-Guideline “Guideline on bioanalytical method validation (http://www.ema.europa.eu/ema/index.jsp?curl=pages/includes/document/document_detail.jsp?webContentId=WC500109686%26mid=WC0b01ac058009a3dc), which implies proof of reproducibility within a given error range. To each sample, 80 mg glass beads were added. Samples were homogenized using a Precellys24 (PeqLab Biotechnology, Erlangen, Germany) at 4 °C for two times over 25 seconds at 5500 rpm and centrifuged at 4 °C and 10000 x *g* for 5 minutes. 10 µL of the supernatants were directly applied to the assay.

Sample handling was performed by a Hamilton Microlab STAR^TM^ robot (Hamilton Bonaduz AG, Bonaduz, Switzerland) and a Ultravap nitrogen evaporator (Porvair Sciences, Leatherhead, U.K.), beside standard laboratory equipment. Mass spectrometric analyses were done on an API 4000 triple quadrupole system (Sciex Deutschland GmbH, Darmstadt, Germany) equipped with a 1200 Series HPLC (Agilent Technologies Deutschland GmbH, Böblingen, Germany) and a HTC PAL auto sampler (CTC Analytics, Zwingen, Switzerland) controlled by the software Analyst 1.6.2. Data evaluation for quantification of metabolite concentrations and quality assessment was performed with the software MultiQuant 3.0.1 (Sciex) and the Met*IDQ*™ software package, which is an integral part of the Absolute*IDQ*™ Kit. Metabolite concentrations were calculated using internal standards and reported in µM.

### Aspartate and pyruvate supplementation

Aspartate medium was freshly prepared for each experiment. For this purpose 10 mM aspartate (Sigma Aldrich, Taufkirchen, Germany) was dissolved in DMEM High Glucose medium. To avoid effects on the cells through an acidification of the medium by aspartate, the pH of the medium was adjusted to 7.5 using 5 M NaOH. The medium was then sterile filtered with sterile, non-pyrogenic, hydrophilic filters (VWR) and 10 % FCS was added. 1 mM Pyruvate was dissolved in medium. Pyruvate-supplemented medium was then sterile filtered and added to the cells.

### Metformin treatment

An optimal non-toxic metformin concentration was titrated for each cell line (WT MEFs: 5 mM, skin fibroblasts: 4 mM, lung fibroblasts: 4 mM). Metformin was then directly added to the respective aspartate- or pyruvate-containing medium. Cells were incubated with metformin for 72 h. Cell number and proteasome activity after 72 h were used as a read out.

### Lysate preparation for phosphoproteome analysis [39]

Mutator cells (300000 cells per well) were seeded in a 6 well plate the day before treatment. Cells were treated with 10 mM aspartate for 4 h. Pre-chilled (4°C) Sodium deoxycholate (SDC) lysis buffer (4% (w/v) SDC, 100 mM Tris -HCl pH 8.5) was added to the cells and scraped off. To inactivate endogenous proteases and phosphatases and ease lysis the lysates were heated to 95 °C for 5 min. The lysates were homogenized at 4°C with the Bioruptor (Diagenode) (2 cycles at maximum output power). To determine the protein concentration BCA assays were performed and the lysates were diluted to identical protein concentrations of appropriate starting material in a final volume of 270 µL SDC lysis buffer. Reduction of caramidomethylate cysteine residues and disulphide bonds was performed by adding 30 µL of reduction/alkylation buffer in a ratio 1:10 to the lysates and incubation for 5 min, at 45 °C and 1500 rpm. After lysates had been cooled down to RT, the enzymes lys-C (Wako) and trypsin (Sigma) were added (ratio of 1:100) and digestion was carried out overnight at 37 °C with shaking (1500 rpm).

### Phosphopeptide enrichment

C8 StageTips (Supelco) (were prepared for each sample as described in the protocol by Rappsilber et al. [69]. Before adding 100 µL EP enrichment buffer (48% (vol/vol) Trifluoroacetic acid (TFA), 8 mM KH_2_PO_4_) to the samples (mix 1500 rpm, 30 sec), 400 µL isopropanol was added and mixed for 30 sec, 1500 rpm to prevent precipitation. The ratio of TiO_2_ beads to protein should be 12:1 (wt:wt). The beads were resuspended in EP loading buffer (6% (vol/vol) TFA/80% (vol/vol) acetonitrile (ACN)) at a concentration of 1 mg/µL and an aliquot of suspended beads was pipetted to the samples and the samples were shaken for 5 min, 40 °C, 2000 rpm. The beads were pelleted via centrifugation at 2000g for 1 min. The supernatant (the non-phosphopeptides) was discarded. The pellets were the washed 4x with 1 mL EP wash buffer (5% (vol/vol) TFA/60% isopropanol (vol/vol)) by shaking at RT for 30 sec, 2000 rpm. After each washing step the beads were pelleted (2000g, 1 min) and the supernatant was discarded. After the final washing step the beads were resuspended in 75 µL EP transfer buffer (0.1 % TFA/60 % (vol/vol) isopropanol) and transferred onto the top of a C8 StageTip. To ensure all beads were transferred from the vial another 75 mL EP transfer buffer were added to each sample and transferred to the C8 StageTip. The StageTips were then centrifuged at 1500 g for 8 min at RT to dryness to pack the TiO_2_ beads on the top of the C8 material. Elution of the phosphopeptides was conducted with 30 µL EP elution buffer (200 μl of ammonia solution (NH_4_OH) added to 800 μl of 40% (vol/vol) ACNacetonitril) and centrifugation at 1500g, 4 min at RT to dryness. The eluates were collected into clean PCR strip tubes and elution was repeated with 30 µL EP elution buffer into the same tubes. The tubes were then placed into an evaporative concentrator and concentrated under vacuum at 45 °C until ∼ 15 µL sample remained.

### StageTip desalting of phosphopeptides

SDB-RPS StageTips (Supelco) were prepared for each sample as described in the protocol by Rappsilber et al [69]. 100 µL of SDB-RPS loading buffer (1% (vol/vol) TFA in isopropanol) were added to each sample and then transferred onto the top of a SDB-RPS StageTip. Centrifugation was carried out at 1500g for 8 min, RT until dryness using the adapter for the StageTips. StageTips were washed with SDB-RPS wash buffer 1 (1% (vol/vol) TFA in isopropanol) and centrifuged until dryness at 1500g, 8 min, RT. Next, one washing step with SDB-RPS wash buffer 2 (0.2% (vol/vol) TFA/5% (vol/vol) ACN) was performed. By addition of 60 µL SDB-RPS elution buffer (Add 20 μl of ammonia solution (NH_4_OH) to 4 mL of 60% (vol/vol) ACNl) the phosphopeptides were eluted (1500g, 5 min, RT) and collected into clean PCR strip tubes. Under vacuum at 45 °C the samples were dried in an evaporative concentrator. Samples were resuspended in 7 µL MS loading buffer (0.1% TFA/2% (vol/vol) ACN) by incubation in a sonicating water bath on low power for 5 min. Afterwards phophopeptides were spun down at 2000g for 1 min at RT. Detection of phosphopeptides via Mass spectrometry was performed according to the EasyPhos protocol.

### Filter-aided sample preparation for proteomic analysis

Protein content was determined using the Pierce BCA protein assay (Thermo Scientific, Rockford, USA) and 10 µg RIPA protein lysate (50 mM Tris, 150 mM NaCl, 1 % NP-40, 0.5 % Sodium deoxycholate, 0.1 % SDS) was subjected to tryptic digest using a modified filter-aided sample preparation (FASP) protocol [70, 71]. Peptides were collected by centrifugation and acidified with 2 µl 100 % trifluoroacetic acid prior to mass spectrometric analysis.

### Proteomic analysis

For LC-MS/MS acquisition a QExactive HF mass spectrometer (ThermoFisher Scientific, Dreieich, Germany) coupled to a RSLC system (ThermoFisher Scientific) was used. Approx. 0.5 µg of peptides were automatically loaded to the HPLC system equipped with a nano trap column (300 μm inner diameter × 5 mm, packed with Acclaim PepMap100 C18, 5 μm, 100 Å; LC Packings, Sunnyvale, CA). After 5 min, peptides were eluted from the trap column and separated using reversed phase chromatography (Acquity UPLC M-Class HSS T3 Column, 1.8 µm, 75 µm x 250 mm, Waters, Milford, MA) by a nonlinear gradient of 0.1 % formic acid in acetonitrile ranging from 3 % to 41 %. The high-resolution (60 000 full width at half-maximum) MS spectrum was acquired with a mass range from 300 to 1500 m/z with automatic gain control target set to 3 x 10^6^ and a maximum of 50 ms injection time. From the MS prescan, the 10 most abundant peptide ions were selected for fragmentation (MSMS) if at least doubly charged, with a dynamic exclusion of 30 seconds. MSMS spectra were recorded at 15 000 resolution with automatic gain control target set to 1 x 10^5^ and a maximum of 100 ms injection time. Normalized collision energy was set to 28 and all spectra were recorded in profile type.

### Progenesis QI for label-free quantification

Spectra were analyzed using Progenesis QI software for proteomics (Version 3.0, Nonlinear Dynamics, Waters, Newcastle upon Tyne, U.K.) for label-free quantification as previously described [71] with the following changes: spectra were searched against the Swissprot mouse database (Release 2017.02, 16686 sequences). Search parameters used were 10 ppm peptide mass tolerance and 20 mmu fragment mass tolerance. Carbamidomethylation of cysteine was set as fixed modification and oxidation of methionine and deamidation of asparagine and glutamine was allowed as variable modification, with only one missed cleavage site. Mascot integrated decoy database search was set to a false discovery rate (FDR) of 1 % with a percolator ion score cut-off of 13 and an appropriate significance threshold p.

### Mitominer analysis

Mitochondrial assignment of proteins was assessed using the Mitominer software (Mitominer 4.0) [72]. Analysis was performed based on gene symbols.

### Bioinfomatic MS data analysis

Progenesis output tables were analyzed using the Perseus software suite (version 1.5.8.7) [73]. Log2 transformed mass spectrometry intensity values were filtered to have at least three out of four quantified values in either the WT or the mutator group. Zero values were imputed with a normal distribution of artificial values generated at 1.6 standard deviations, subtracted from the mean, of the total intensity distribution and a width of 0.3 standard deviations. This places the imputed values at the lower limit of the intensity scale, which represents detection limit of the used instrumentation. For gene annotation enrichment analysis of the data from isolated mitochondria, we used 710 proteins that were confirmed to be true mitochondrial proteins based on the Mitominer software (see above).

Gene annotation enrichment analysis was performed with the 1D annotation enrichment algorithm as previously described [74]. As gene annotations for significance tests, we used the Uniprot Keyword annotation as well as Gene Ontology terms Biological process (GO:BP), Molecular function (GO:MF) and Cellular Component (GO:CC) [75]. In brief, it is tested for every annotation term whether the corresponding numerical values have a preference to be systematically larger or smaller than the global distribution of the values for all proteins, which is reported as normalized enrichment score. Additional pathway analysis was performed with the DAVID Bioinformatic Resources 6.8.

### Cell proliferation assay

20000 cells per cell line were seeded as triplicates in 6 well plates. After attachment of the cells overnight control wells were counted for each cell line to determine the starting cell number after cell seeding. For aspartate supplementation cells were washed with PBS and 4 ml DMEM High Glucose medium containing 10 mM aspartate was added to the wells. Cells were then counted again at day 5 of the experiment to determine the final cell number and doubling rates per day were calculated as previously described by Sullivan et al. [8].

### Protein synthesis assay (Aspartate treatment of mutator MEFs)

20000 cell per mutator MEF cell line (n=4) were seeded on coverslips in 6 well plates. After overnight regeneration cells were treated with 10 mM aspartate supplemented in high glucose medium for 48 h. Cells were then treated for 4 h with control medium or with medium containing 100 µM cycloheximide. After the treatment, medium was removed and cells were incubated with growth medium containing EZClick^TM^ O-propargyl-puromycin (OPP) reagent (Biovision, Milpitas, USA) for 30 min. Afterwards, OPP-containing medium was removed, cells were washed with PBS and fixed with 4 % PFA for 15 min. After fixation, cells were permeabilized in 0.5 % TritonX-100 in PBS for 15 min, washed twice with PBS and stained by adding 500 µl EZClick^TM^ fluorescence azide reaction cocktail for 30 min. Afterwards, cells were washed with PBS and nuclei were stained by incubation with 4’-6-Diamidin-2-phenylindol (DAPI) (Sigma-Aldrich) in PBS for 30 min. Cells were washed again with PBS and mounted on object slides with DAKO mounting medium (DAKO, Hamburg, Germany). Fluorescence was measured using the LSM710 fluorescence microscope (Zeiss).

### siRNA mediated mRNA silencing

We first determined the optimal siRNA concentrations for efficient knockdown of p27, p28 and Raptor (10 nM) which did not affect cell viability and proliferation. As Rpn6 is an essential proteasome subunit, we performed only a partial knockdown of this subunit (0.3 nM), which did not affect cell growth. Reverse transfection of mutator MEFs was used to deliver siRNAs. Two different siRNAs and control siRNAs were used for each of the different targets (Psmd10: s203895+s79154; Psmd11: s87416+s87415; Psmd9: s84561+s84562, Control siRNA Negative Control #1+#2, Ambion, Thermo Fisher Scientific). Raptor knockdown was performed with one Raptor siRNA (s92713) and Control siRNA Negative Control #1. siRNAs were first incubated with Opti-MEM for 5 min at RT and 5 µL of transfection reagent (RNAiMax, Thermo Fisher Scientific) was applied per 6 well. siRNAs were incubated with transfection reagent for further 20 min at RT. Cells were harvested in antibiotic-free medium, counted, and plated in 6 well plated. 20000 cells per well were plated for 72 h aspartate treatment and 100000 cells per well for 48 h of silencing without aspartate. Opti-MEM containing siRNAs and transfection reagents was added to the cells. The next morning medium was changed either with normal DMEM High glucose medium or medium containing 10 mM aspartate. Silencing efficiency was confirmed via immunoblotting of the respective proteins.

### Rapamycin and aspartate cotreatment of mutator cells

Optimal concentration for rapamycin was first determined for mutator cells by assaying phosphorylation of p70S6 kinase. At a concentration of 0.5 nM mTORC1 was specifically inhibited while mTORC2 was not affected (no decrease in the phosphorylation of Akt). Mutator cells were seeded the day before the treatment. 40000 cells per well were used per 6 well. After overnight attachment cells were treated with 4 mL of medium containing 10 mM aspartate and 0.5 nM rapamycin for 72 h. Efficiency of mTOR inhibition was confirmed by reduced phosphorylation of p70S6 kinase.

### Bulk mRNA sequencing

Strand specific, polyA-enriched RNA sequencing was performed as described earlier [76]. Briefly, RNA was isolated from whole-cell lysates using the Total RNA kit (Peqlab, VWR) and RNA integrity number (RIN) was determined with the Agilent 2100 BioAnalyzer (RNA 6000 Nano Kit, Agilent). For library preparation, 1 μg of RNA was poly(A) selected, fragmented, and reverse transcribed with the Elute, Prime, Fragment Mix (Illumina). A-tailing, adaptor ligation, and library enrichment were performed as described in the TruSeq Stranded mRNA Sample Prep Guide (Illumina). RNA libraries were assessed for quality and quantity with the Agilent 2100 BioAnalyzer and the Quant-iT PicoGreen dsDNA Assay Kit (Life Technologies). RNA libraries were sequenced as 150 bp paired-end runs on an Illumina HiSeq4000 platform. The STAR aligner (v 2.4.2a) with modified parameter settings (--twopassMode=Basic) was used for split-read alignment against the human genome assembly hg19 (GRCh37) and UCSC known gene annotation. To quantify the number of reads mapping to annotated genes we used HTseq-count° (v0.6.0) [77]. FPKM (Fragments Per Kilobase of transcript per Million fragments mapped) values were calculated using custom scripts. Normalized data set was analyzed using the Perseus Software as described in the proteomics section.

### Statistical analysis

Data are depicted in the figures as mean ± SEM. All statistical analysis was performed using GraphPad Prism software (Version 7, GraphPad, San Diego, CA, USA). Statistical tests were applied to all experiments where applicable. Student’s unpaired t-test was mainly used to determine significance between WT and Mut MEFs. Student’s paired t-test was used to determine significant differences when different mutator cell lines were treated with aspartate. The one sample t-test was used to determine significance when native gel immunoblotting was performed with one single cell line or different mutator cell lines to eliminate signal intensity differences between replicates or individual mutator cell lines. Significance was indicated in the figures as *: *p*<0.05, **: *p*<0.01 or ***: *p*< 0.001. Statistical analysis of the respective experiments was performed as explained in the figure legends.

## Supplementary Tables

Table S1 Bulk mRNA sequencing.xlsx

Table S2 1D enrichment analysis cell lysates.xlsx

Table S3 1D enrichment analysis isolated mitochondria.xlsx

Table S4 Mass spectrometry data from isolated mitochondria.xlsx

Table S5 Phosphoproteomics.xlsx

Table S6 Phosphoproteomics_Fisher Exact Test using 233 t test sign.xlsx

Table S7 Mass spectrometry data from cell lysates.xlsx

