## Supplemental information for "Adaptive mitochondrial regulation of the proteasome"

### Figure S1: related to Figure 1

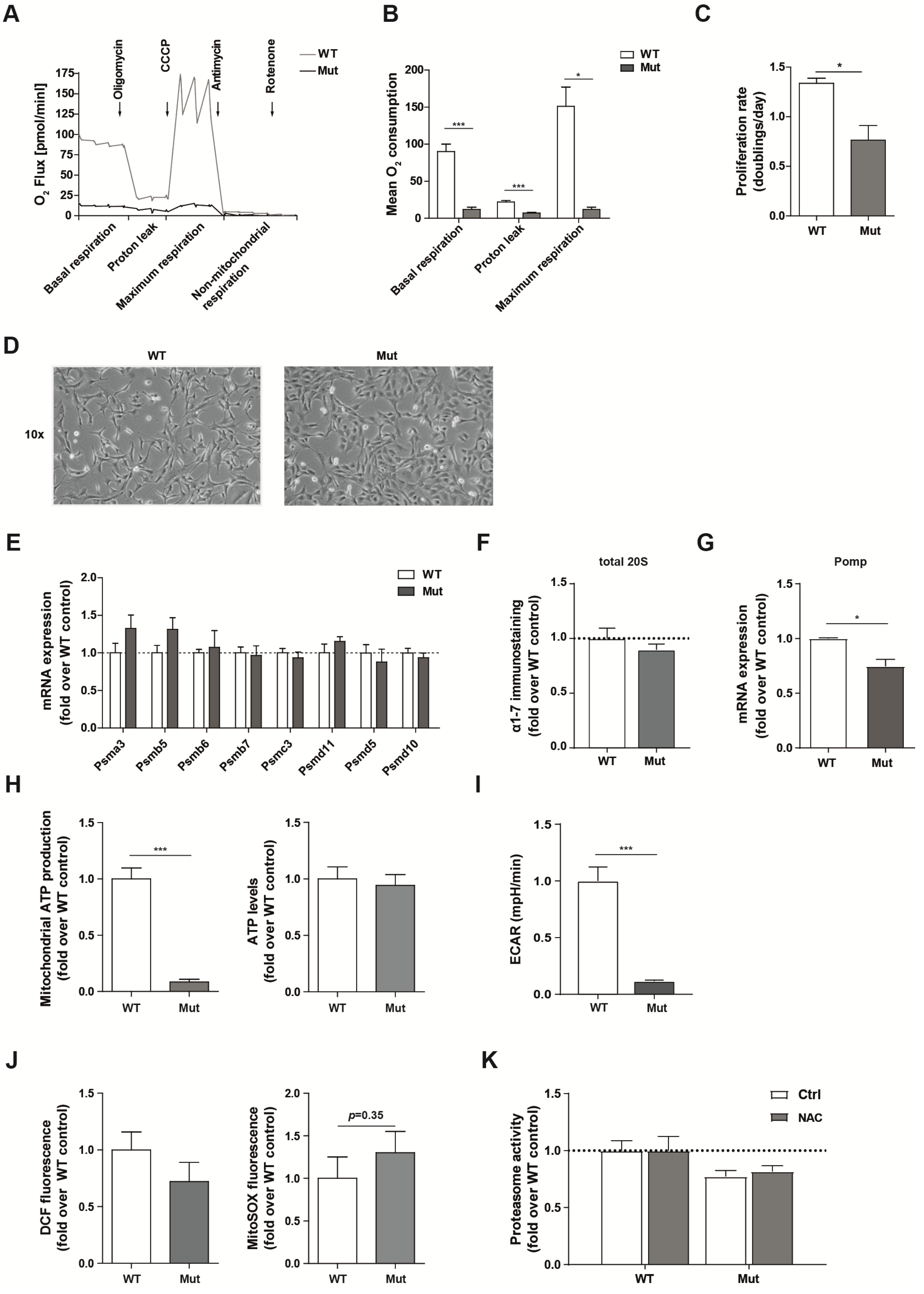

**Figure S2: related to Figure 2**

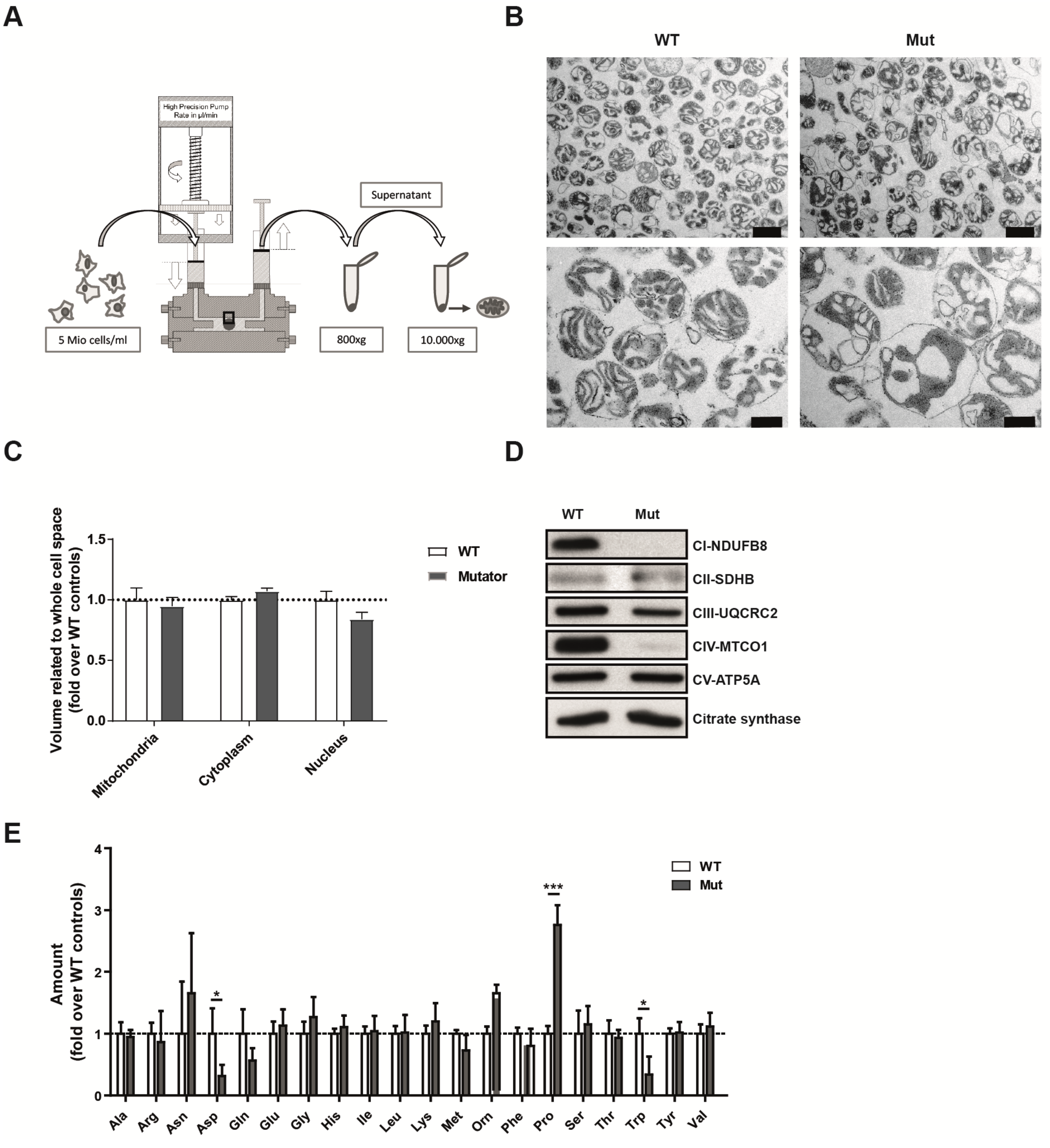

**Figure S3: related to Figure 3**

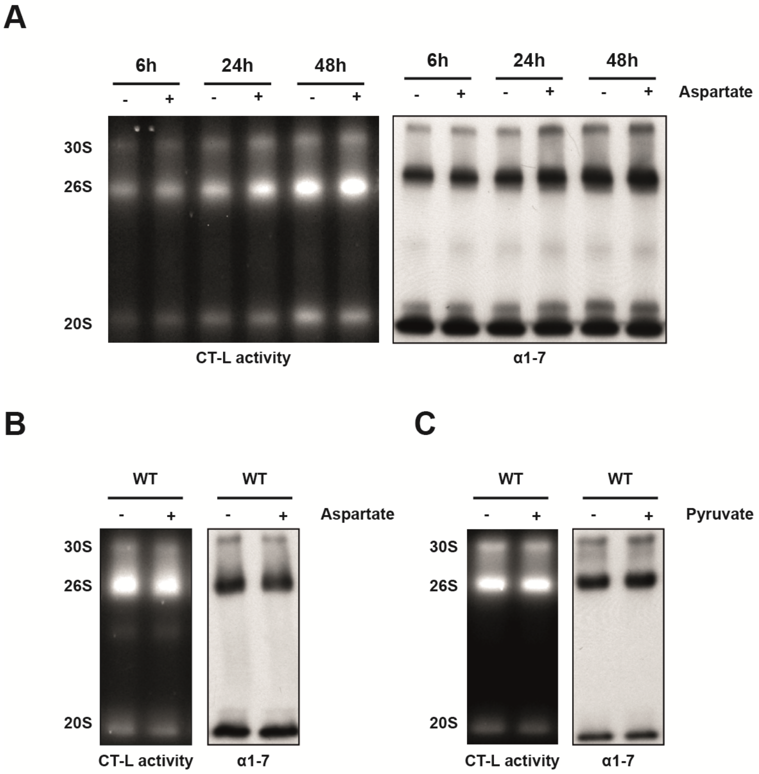

**Figure S4: related to Figure 4**

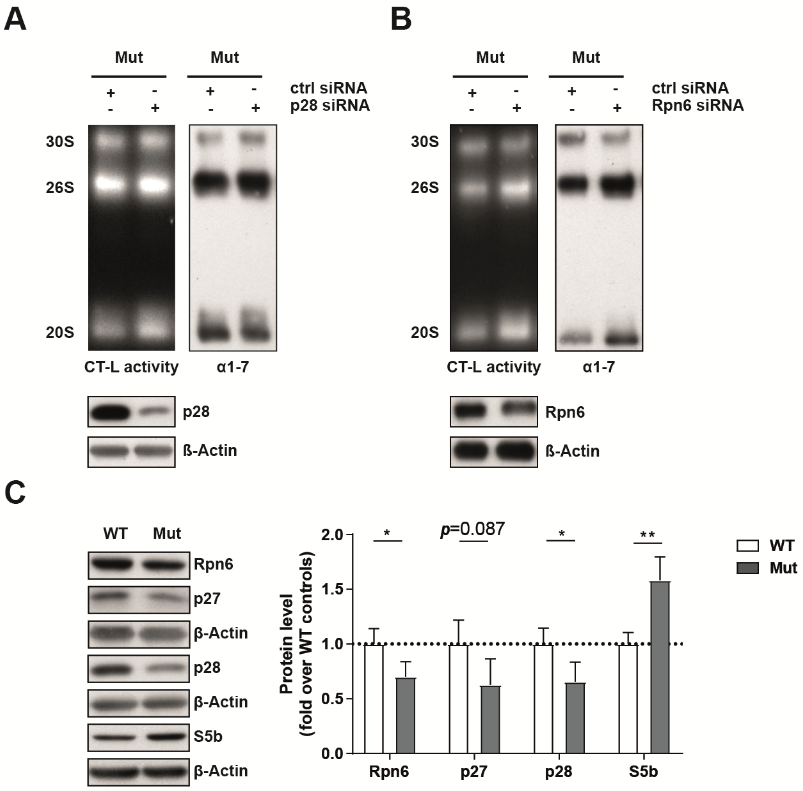

**Figure S5: related to Figure 5**

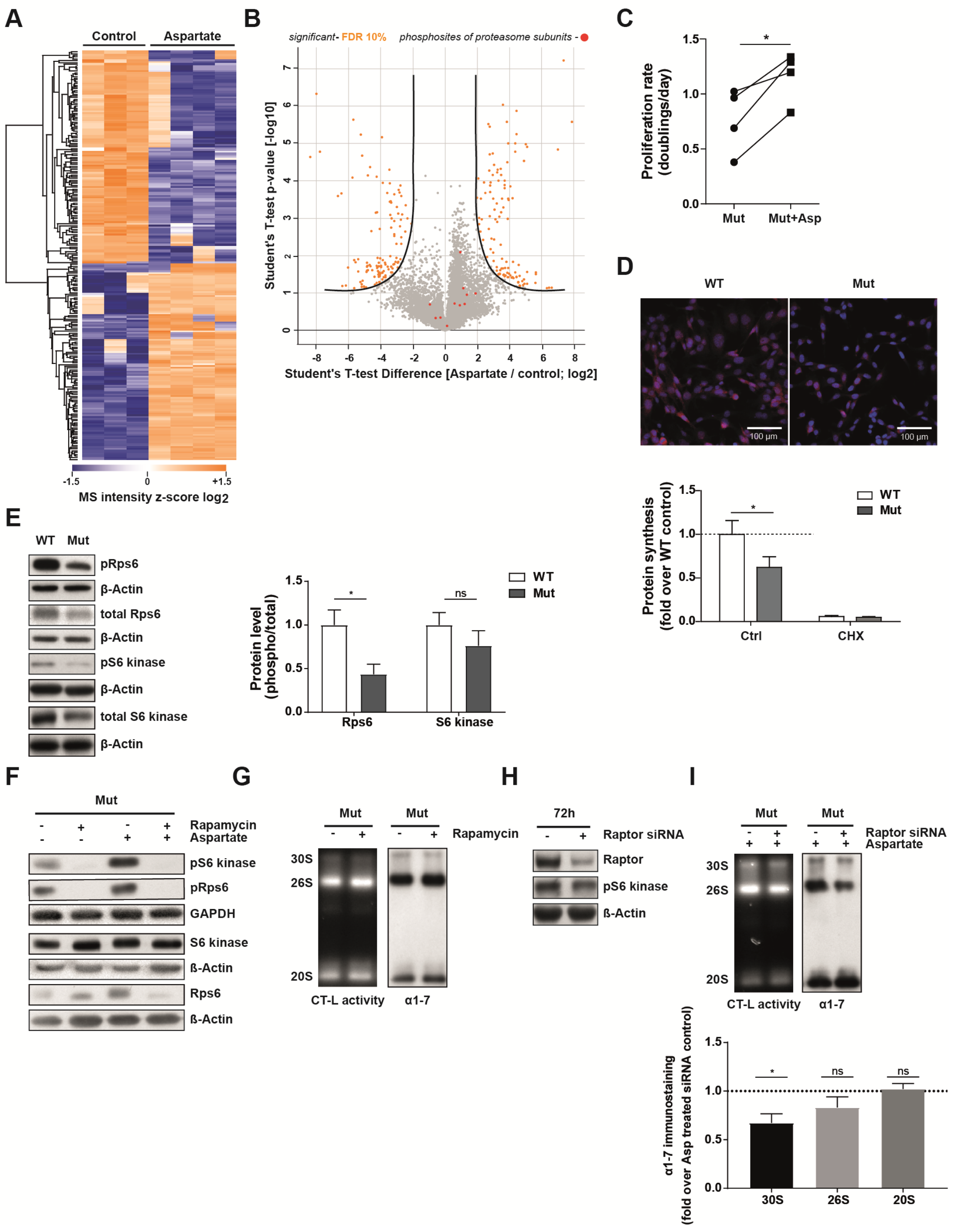

**Figure S5: related to Figure 6**

**A**

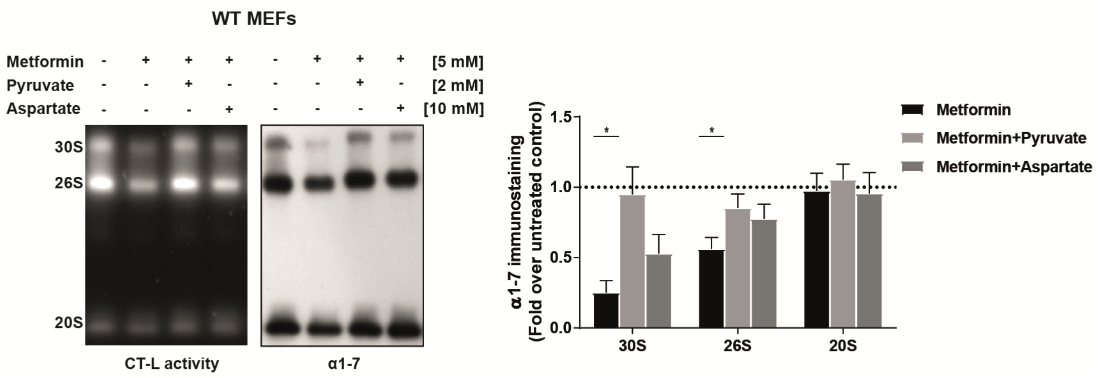


### Supplemental Figure titles and legends

**Figure S1: related to Figure 1**

(A) Mitochondrial oxygen flux monitored over time at base line condition, after addition of oligomycin (determining proton leak), CCCP (assessing maximal respiration), or antimycin A and rotenone (assaying non-mitochondrial respiration) as analyzed by the Seahorse XF analyzer. Curves show mean values from three independent wildtype (WT) and four independent mutator (Mut) cell lines. (B) Mean oxygen consumption rates in WT (n=3) and mutator (n=4) cells. Bar graphs show mean±SEM. Significance was determined using two-way ANOVA with Bonferroni multiple comparison test. (C) Proliferation rates of WT (n=3) and mutator (n=4) cell lines were determined by counting cells at day 1 and day 5 after seeding of the cells. Doublings per day were then calculated as explained in the methods part. Bar graphs show mean±SEM. Significance was determined using student's unpaired t-test. (D) Representative images show cellular morphology of WT and mutator MEFs. Magnification: 10x. (E) RT-qPCR analysis of proteasome subunit mRNA expression in WT (n=3) and mutator (n=4) cells. Data represent mean±SEM relative to WT control. Statistical test: two-way ANOVA with Bonferroni multiple comparison test. (F) Quantification of amounts of total 20S complexes in WT and mutator cells as resolved by blotting of native gels and immunostaining for the 20S subunits α1-7. Bar graph shows combined signals for 30S, 26S and 20S related to WT controls. Significance was determined using student's unpaired t-test. (G) RT-qPCR analysis of Pomp mRNA expression in WT (n=3) and mutator (n=4) cells. Data represent mean±SEM relative to WT control. Statistical test: unpaired t-test. (H) Relative mitochondrial ATP production (oligomycin sensitive respiration) in WT (n=3) and mutator (n=4) cells as analyzed by the Seahorse XF analyzer. Relative cellular ATP levels in WT (n=3) and mutator (n=4) cells as measured by a luciferase-based assay. (I) Extracellular acidification rate (ECAR) was measured using the Seahorse XF analyzer to determine the spare glycolytic capacity in WT (n=3) and Mut MEFs (n=4). Values were normalized to the mean of WT controls. Significance was determined using student's unpaired t-test. (J) Cellular ROS was measured using $H_2$DCFDA staining and flow cytometry in WT (n=2) and mutator (n=2) cell lines. Flow cytometry analysis of WT (n=3) and mutator (n=4) cell lines stained with MitoSOX Red for detection of mitochondrial superoxides. Data represent mean±SEM relative to WT. Significance was determined using student's unpaired t-test. (K) CT-L proteasome activity in WT (n=3) and mtDNA mutator (n=4) MEFsafter treatment with control or 5 mM NAC-containing medium for 2 h. Bars show mean+SEM. Values were displayed as fold change relative to WT.

**Figure S2: related to Figure 2**

(A) Scheme of the mitochondrial isolation procedure. A high precision pump ensures, via gastight syringes, the continuous sample delivery in a constant rate to the "Balch-homogenizer". Cell breakage occurs upon passage through a defined clearance (square), which is adjusted by selecting tungsten carbide balls of different diameters. The cell homogenate is collected in a second syringe and transferred to a 2ml Eppendorf cup for differential centrifugation. (B) Representative electron microscopy images of mitochondria isolated from one WT and mutator cell line. Scale bar: upper panel 1 µm, lower panel 500 nm. (C) Proportion of mitochondrial volume was quantified as described by Hacker and Lucocq (Hacker and Lucocq, 2014). In total, 30 electron micrographs (1000x magnification) from three wildtype clones (two technical replicates, each) and 43 electron micrographs (1000x magnification) from four mutator clones (two technical each) were used for quantification. Data represent mean±SEM relative to WT controls. Statistical test: student's unpaired t-test. (D) Representative Western blot analysis of single respiratory chain complex subunits in mitochondria isolated from one WT and mutator cell line. Four independent isolations were used for Western blots. Citrate synthase served as a loading control. (E) Quantification of the amount of 20 amino acids in WT (n=3) and mutator (n=4) cells. 6 replicates were measured for each cell line and the respective values were normalized to the cell number of each cell line. Data represent mean±SEM relative to WT controls. Statistical test: student's unpaired t-test.

**Figure S3: related to Figure 3**

(A) Representative native gel analysis of active proteasome complexes in cell lysates from one mutator cell line treated with 10 mM aspartate for 6 h, 24 h and 48 h. Chymotrypsin-like (CT-L) substrate overlay assay and immunoblotting for 20S α 1-7 subunits is shown. (B+C) Native gel of active proteasome complexes in cell lysates from WT cells (n=3) treated with 10 mM aspartate or 1 mM Pyruvate for 72 h. Chymotrypsin-like (CT-L) substrate overlay assay and immunoblotting for 20S α 1-7 subunits is shown.

**Figure S4: related to Figure 4**

(A+B) Representative native gel analysis of active proteasome complexes in cell lysates from one mutator cell line upon Rpn6 (n=5) and p28 (n=4) silencing for 72 h with chymotrypsin-like (CT-L) substrate overlay assay, immunoblotting for 20S α 1-7 and quantification thereof. Control cells were transfected with a combination of two different control siRNAs. (C) Western blot analysis of 26S proteasome assembly factor expression in WT and mutator cells. β-Actin was used as a loading control. Densitometric analysis shows mean±SEM from three WT and four mutator cell lines. Significance was determined using student's unpaired t-test.

**Figure S5: related to Figure 5**

(A) Heatmap of 233 phosphosites significantly regulated by aspartate treatment compared to non-treated controls. Each row corresponds to a single distinct phosphosite. Rows are ordered according to unsupervised hierarchical clustering (Pearson correlation of z-score), after imputation to replace missing values. (B) The depicted volcano plot shows significantly altered phosphorylation sites relative to controls with a 10 % FDR. Phosphosites of proteasome subunits are shown in red. (C) Proliferation rates of mutator (n=4) cell lines treated with 10 mM aspartate were determined by counting cells at day 1 and day 5 after seeding of the cells. Doublings per day were then calculated as explained in the methods part. Graph shows increase for each mutator cell line after aspartate supplementation. Significance was determined using student's paired t-test. (D) Representative fluorescence images showing nascent protein synthesis (red signal) and cell nuclei (blue signal) in WT (n=3) and mutator (n=4) cells. (Cycloheximide CHX) treatment with 100 µM for 4 h served as a negative control. Scale bar: 100 µm. Graph shows mean±SEM relative to WT for Ctrl and CHX treated samples. Statistical test: Two-way ANOVA with Bonferroni multiple comparison test. (E) Analysis of mTOR signaling in WT (n=3) mutator (n=4) cells. Representative Western blots of total and phosphorylated levels of p70 S6 kinase and S6 ribosomal protein (Rps6). Bar graphs show β-Actin normalized phospho-protein levels related to total levels of the respective protein in WT and mutator cells (Mean±SEM). Significance was determined using student's unpaired t-test. (F) Analysis of mTOR signaling upon treatment with 0.5 nM rapamycin and 10 mM aspartate for 72 h in one mutator cell line. Representative Western blots of total and phosphorylated levels of p70 S6 kinase, S6 ribosomal protein (Rps6). GAPDH and β-Actin was used as a loading control. (G) Representative native gel analysis of active proteasome complexes in cell lysates from one mutator cell line upon rapamycin treatment (0.5 nM) for 72 h. Chymotrypsin-like (CT-L) substrate overlay assay and immunoblotting for 20S α 1-7 subunits is shown (H) Protein levels of Raptor and phospho S6 kinase after aspartate supplementation and siRNA mediated Raptor silencing in one mutator cell line (n=4 independent experiments). (I) Representative native gel analysis of active proteasome complexes in cell lysates from one mutator cell line upon 72 h treatment with 10 mM aspartate and siRNA mediated Raptor silencing. Chymotrypsin-like (CT-L) substrate overlay assay and immunoblotting for 20S α 1-7 subunits is shown. Densitometric analysis shows mean±SEM of fold change over control from four independent experiments. Significance was determined using the one sample t-test.

**Figure S6: related to Figure 6**

Representative native gel analysis of active proteasome complexes in cell lysates from one WT MEF cell line cotreated with 10 mM aspartate or 1 mM pyruvate together with 5 mM metformin for 72 h. Chymotrypsin-like (CT-L) substrate overlay assay and immunoblotting for 20S $\alpha$ 1-7 subunits is shown. Bar graphs represent mean±SEM relative to respective untreated WT MEF cell line (n=4 independent experiments). Significance was determined using one sample t-test.
